# Evidence of post-domestication hybridization and adaptive introgression in Western European grapevine varieties

**DOI:** 10.1101/2021.03.03.432021

**Authors:** S. Freitas, M.A. Gazda, M. Rebelo, A.J. Muñoz-Pajares, C. Vila-Viçosa, A. Muñoz-Mérida, L.M. Gonçalves, D. Azevedo-Silva, S. Afonso, I. Castro, P.H. Castro, M. Sottomayor, A. Beja-Pereira, J. Tereso, N. Ferrand, E. Gonçalves, A. Martins, M. Carneiro, H. Azevedo

## Abstract

Grapevine (*Vitis vinifera* L.) is one of the most significant crops in the world. Today’s richness in grapevine diversity results from a complex domestication history over multiple historical periods. Here, we employed whole genome resequencing to elucidate different aspects of the recent evolutionary history of this crop. Our results support a model in which a central domestication event in grapevine was followed by post-domestication hybridization with local wild genotypes, leading to the presence of an introgression signature in modern wine varieties across Western Europe. The strongest signal was associated with a subset of Iberian grapevine varieties, which show large introgression tracts. We targeted this study group for further analysis, demonstrating how regions under selection in wild populations from the Iberian Peninsula were preferentially passed on to the cultivated varieties by geneflow. Examination of underlying genes suggests that environmental adaptation played a fundamental role in both the evolution of wild genotypes and the outcome of hybridization with cultivated varieties, supporting a case of adaptive introgression in grapevine.

## INTRODUCTION

The European grapevine (*Vitis vinifera* L.) is one of the most charismatic plants of the Mediterranean agricultural landscape, and one of the most widely grown fruit crops in the world. *V. vinifera* has diverged into two subspecies that distinguish cultivated varieties (*V. vinifera* ssp. *vinifera*) from their wild relatives (*V. vinifera* ssp. *sylvestris*), hereafter *vinifera* and *sylvestris*, respectively (1). *V. vinifera* is the only species of the *Vitis* genus’ European clade, holding wild populations that occupy river banks and damp woods from the Transcaucasian region of West Asia to the Iberian Peninsula (2, 3). The origin and domestication centers for *V. vinifera* are both widely considered to match the Transcaucasian region, as suggested by morphological data and the presence of higher nucleotide diversity in both wild and cultivated germplasm belonging to this region (4–6). The earliest archaeological evidence for domestication of Eurasian grapevine was observed in the South Caucasus, timing it to the Late Neolithic Period, 8000 years ago (ya) (7). Divergence between the two subspecies, estimated using whole genome resequencing, places this event much earlier (22-400 kya) with an important bottleneck occurring *ca.* 8000 ya (8, 9). Between 6000 and 3000 ya, viticulture spread south and west across the Mediterranean space (10).

Like most crops, grapevine still retains numerous unanswered questions regarding its origin and domestication path (11). A restricted origin hypothesis first predicted that diversity of cultivated grapes was limited to a few founder genotypes in a single location (12). Since then, use of molecular markers spanning a restricted number of loci has suggested the existence of gene flow between cultivated varieties and local wild genotypes, particularly associated with Western European wine varieties. Thus, there has been conflicting molecular evidence to support either multiple independent domestications (4, 13), or an initial domestication in the Transcaucasus followed by post-domestication hybridization with local wild relatives (14). Characterization of cultivated and wild genetic diversity across the Mediterranean geographic range has placed the Iberian Peninsula region as a strong candidate for local contribution of wild populations to the overall modern-day structure of the cultivated grape (4). The Iberian Peninsula is a hotspot for grapevine genetic diversity supporting hundreds of autochthonous varieties. Its diversity has been structured into three genetic groups that reflect either a core Iberian nature, a common genetic signature with Western and Central Europe varieties, or a Western Mediterranean provenance that incorporates the Maghreb (15). This complexity reflects an ancient and rich domestication history, which translates into the diversity and identity of Iberian wines like the distinctive Vinhos Verdes.

Since the earliest domestication period, human activity led to the creation of thousands of grapevine varieties (1). Whole genome resequencing currently offers new opportunities to address this domestication path. Here, we analyzed 100 whole-genome sequences from the *Vitis* genus, to elucidate important aspects of grapevine domestication, including the singularity of Western European genotypes in their relationship with sympatric *sylvestris* genotypes. Our data supports the hypothesis of a single major domestication event in *Vitis vinifera*, followed by post-domestication hybridization with wild relatives in Western Europe. A subset of Iberian varieties exhibited particularly extensive signs of unidirectional introgression with local wild genotypes. Analysis of introgression tracts and selective sweep mapping suggest that regions under selection in donor wild genotypes were favorably incorporated into introgressed regions of the Iberian varieties. Ensuing analysis of the underlying genes suggests a strong association with environmental adaptation, and a scenario of adaptive introgression as the result of hybridization with local wild relatives.

## RESULTS

### Whole-genome resequencing dataset

In *V. vinifera*, several reports have emphasized the importance of Western Europe as a source of either an independent domestication, or of post-domestication hybridization between cultivated forms and local wild plants (4, 6, 14). To provide a detailed understanding of this past genetic history, we collected 51 cultivated (*vinifera*) samples, representing a large geographic range reinforced with Western European and particularly Iberian varieties, nine wild (*sylvestris*) samples from the Iberian Peninsula, and three *Vitis* sp. genotypes (Table S1). Genomes were resequenced using Illumina technology. This dataset was complemented with publicly available sequencing data from 37 other genotypes (Table S1). The total dataset comprised 100 genotypes, and was composed of several *Vitis sp.* members, *sylvestris* samples from eastern and Iberian origins, and an array of table and wine *vinifera* varieties. Reads were mapped to the *V. vinifera* reference genome (Pinot Noir PN40024). Samples mapped on average to 86% of the genome, regardless of origin, showing a mapping depth of 5.2X (Fig. S1).

### Patterns of population structure

We began by investigating population structure between species within the *Vitis* genus and cultivated and wild *V. vinifera* samples, using clustering methods based on genotype likelihoods (16–18). First, principal components analysis (PCA) was used to help visualize the relationships between the different *Vitis* species members (Fig. 1A; Fig. S2A,B). *Vitis rotundifolia* (also designated *Muscadinia rotundifolia*) was clearly divergent from the remaining genotypes, in line with previous WGS assessments (9, 19). Cultivated and wild *V. vinifera* formed a consistent cluster, with wild genotypes appearing closer to remaining *Vitis* species. We then resolved the relationship between cultivated and wild genotypes by excluding non-*V. vinifera* samples (Fig. 1B; Fig. S2C,D). This resulted in a clear separation between wild and cultivated varieties (PC2, 2.88% of the variance). Most significantly, an overall east-to-west geographical gradient was evident both in wild and cultivated samples (PC1, 3.71% of the variance), consistent with a parallel westward expansion in these subspecies. Within cultivated varieties there was a separation between table and wine varieties (Fig. 1B). The latter consisted mostly of genotypes from Western and Central Europe and the Iberian Peninsula. Here, varieties with estimated Iberian provenance were split into two groups, distinctly separated by varieties of Western and Central Europe estimated provenance. Overall results were strongly supported by ancestry and phylogenetic tree analysis (Fig. 1C-D). Subsequently, we used population structure data and the perceived geographical origin of the varieties, to cluster genotypes within six study groups: wild samples were divided between Eastern and Iberian groups (W_EAST_; W_IBERIA_), while cultivated varieties were divided between one table (C_TABLE_) and three wine groups, representing Western and Central Europe (C_wWCE_) and the Iberian Peninsula (C_wIB1_ and C_wIB2_) (Fig. 1B-D). We estimated Identical-by-descent (IBD) scores to single out the presence of clones, in which case, a single clone was retained for subsequent population studies (Fig. S3, Table S2).

**Fig. 1.**
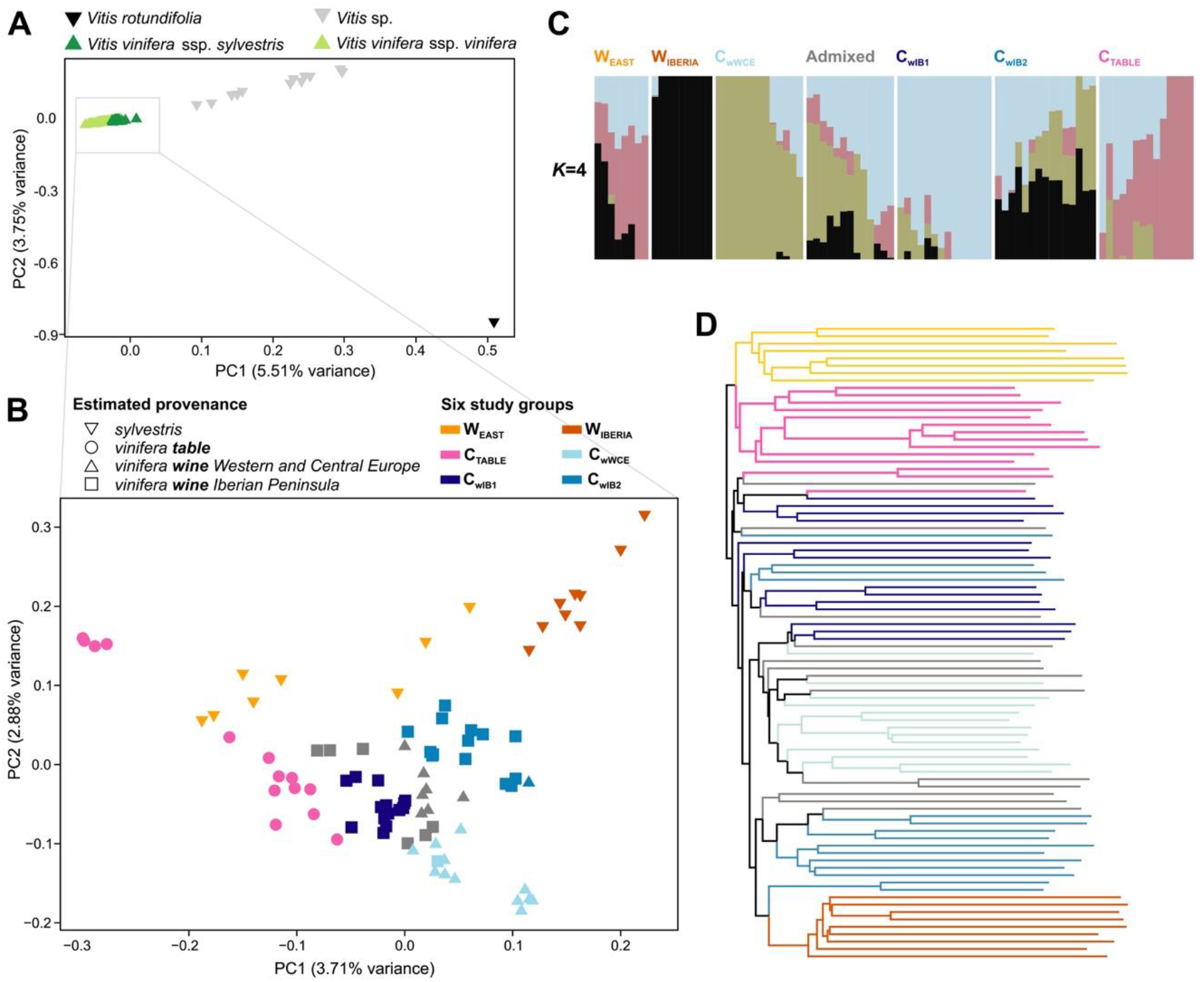
Population structure analysis of *Vitis sp.* and *Vitis vinifera* genotypes. (**A**) Principal components analysis (PCA) plot of *Vitis* sp. and *Vitis vinifera* wild and cultivated genotypes. (**B**) PCA plot of *Vitis vinifera* samples. (**C**) Ancestry proportions of all *Vitis vinifera* genotypes following admixture analysis for *K*=4; bars represent individual genotypes, organized into six study groups plus remaining admixed individuals. (**D**) Phylogenetic tree of *Vitis vinifera* samples.

When interrogating these study groups for patterns of ancestry using admixture analysis, C_wIB2_ displayed compelling evidence of shared ancestry with Iberian *sylvestris* genotypes (Fig. 1C, Fig. S4A). These results were highly contrasting with its C_wIB1_ counterpart in K=4 and other meaningful simulations (K=2 to K=6). Furthermore, C_wIB2_ was the wine group closest to W_IBERIA_ genotypes in the PCA (Fig. 1B), and phylogenetic tree analysis placed most of the C_wIB2_ genotypes within the same subclade as W_IBERIA_ samples (Fig. 1D; Fig. S4B). All this evidence amounts to a high likelihood of genetic relatedness between C_wIB2_ and W_IBERIA_ groups. Additionally, wine varieties from Western and Central Europe were also polarized by PC1 of the multivariate analysis to comparable levels as W_IBERIA_ and C_wIB2_ (Fig. 1B). They also clustered next to these two groups in the phylogenetic tree, suggesting some extent of admixture between C_wWCE_ and W_IBERIA_, though smaller than that observed for C_wIB2_. Table varieties clustered closer to wild genotypes from the center of origin (W _EAST_) in all analysis (Fig. 1), supporting previous evidence of high phenotypic and genetic differentiation between table and wine grapes (9, 20). This also indicates that a potential hybridization event in Western European grapevine varieties took place after differentiation between wine and table grapes.

### Introgression testing between cultivated varieties and local wild genotypes

Given the potential for hybridization with local wild relatives displayed by C_wIB2_, we next tested for admixture using Patterson’s *D* statistics – an explicit test of gene flow (Fig. 2A) (21). We configured the statistic to test potential gene flow from *sylvestris* donor groups (either W_EAST_ or W_IBERIA_), to recipient cultivated groups. When placing C_TABLE_ (which previously showed no evidence of admixture with W_IBERIA_) as P1, wine genotypes as P2, and W_IBERIA_ as the donor group, we observed positive and statistically significant Patterson’s *D* levels across all tests (Fig. 2B). Results were strongest for C_wIB2_ (*D*=0.1772; *Z*=114.2490; *P*=0.0000), followed by C_wWCE_ (*D*=0.1448; *Z*=76.7658; *P*=0.0000) and C_wIB1_ (*D*=0.0927; *Z*=59.4755; *P*=0.0000). Conversely, *D* scores were negative and much closer to zero (neutrality) when replacing W_IBERIA_ with W_EAST_ (Fig. 2B, Table S3). Statistical support for the strongest gene flow occurring between W_IBERIA_ and C_wIB2_ was still evident when strictly wine groups were placed as recipients (Fig. 2C). When looking at each chromosome separately, there was a highly heterogenous distribution of *D* (Fig. 2D), indicating that exchange of genetic information was not random across the genome. We found that chromosomes 6 (D=0.20) and 14 (D=-0.03) provided extreme cases of high and low admixture, respectively. Interestingly, when extending this analysis to mitochondrial and chromosomal genomes, we observed an increase in Patterson’s *D*, which suggests a *sylvestris* maternal progenitor in the cross(es) that resulted in admixture. Collectively, these results support the first genome-level claim of admixture between Western European grapevine varieties and local wild relatives.

**Fig. 2.**
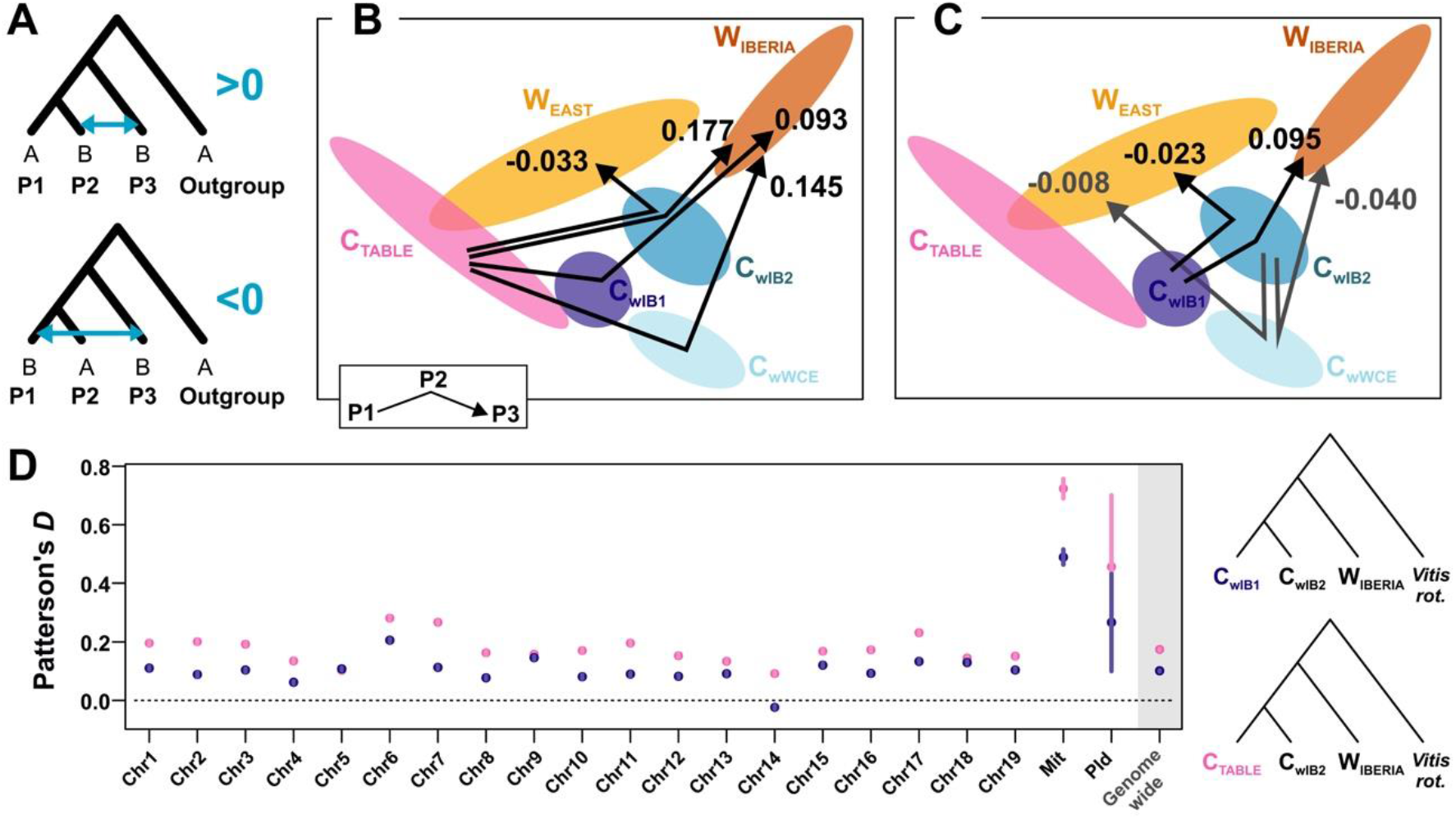
Patterson’s *D* statistics test for admixture. (**A**) Patterson’s *D* statistics (or ABBA-BABA test) assumes that in four groups phylogenetically related as such -(((P1,P2),P3),O) – the proportion of ABBA and BABA sites will be equal under a scenario of incomplete lineage sorting without gene flow. Genome-wide levels of introgression from the donor P3 group can be detected by the presence of statistically significant levels of excess ABBA (P3→P2) or BABA (P3→P1) patterns. (**B**) Genome-wide Patterson’s *D* scores when confronting table (P1) against wine groups as P2, and wild groups as P3. (**C**) Genome-wide Patterson’s *D* scores assuming wine groups as either P1 or P2, and wild groups as P3. (**D**) Chromosome-level estimates of Patterson’s *D* statistics (±s.e.) estimated in the C_TABLE_ or C_wIB1_ (P1), C_wIB2_ (P2) and W_IBERIA_ (P3) configurations.

### Analysis of introgression impact

We next looked at the impact of introgression by performing a genome-wide characterization of three separate properties of the data, 1) nucleotide diversity, 2) IBD scores, and 3) genetic differentiation. Diversity indexes and measures of population differentiation were estimated across the genome while taking into account nucleotide uncertainty (16) (Fig. 3; Fig. S5). Results for nucleotide diversity (*π*) (Fig. 3A) and Watterson’s theta (*θ_W_*) (Fig. S5A) showed a decrease in diversity when comparing wild grapes from the East (*π*=0.0140; *θ_W_*=0.0140) and the wild Iberian group (*π*=0.0112; *θ_W_*=0.0108). There was also a decrease in nucleotide diversity from wild to table (*π*=0.0120) and wine (*π*=0.0110 to 0.0119) groups, which is consistent with the presence of a weak domestication bottleneck (14). Comparisons were all statistically supported (*P*<0.001). Interestingly, there was an increase in nucleotide diversity in C_wIB2_ when compared to the remaining wine groups, which is compatible with higher proportion of admixture following introgression of local *sylvestris* into the *vinifera* genetic pool. Tajima’s D statistics were close to zero for W_EAST_ (TD=−0.0346), but they increased for remaining study groups (Fig. 3B), consistent with population contraction (a likely scenario in the wild Iberian population). An overall similar profile was observed when we summarized IBD scores between all individuals of a study group against all individuals of a second group (Fig. 3C).

**Fig. 3.**
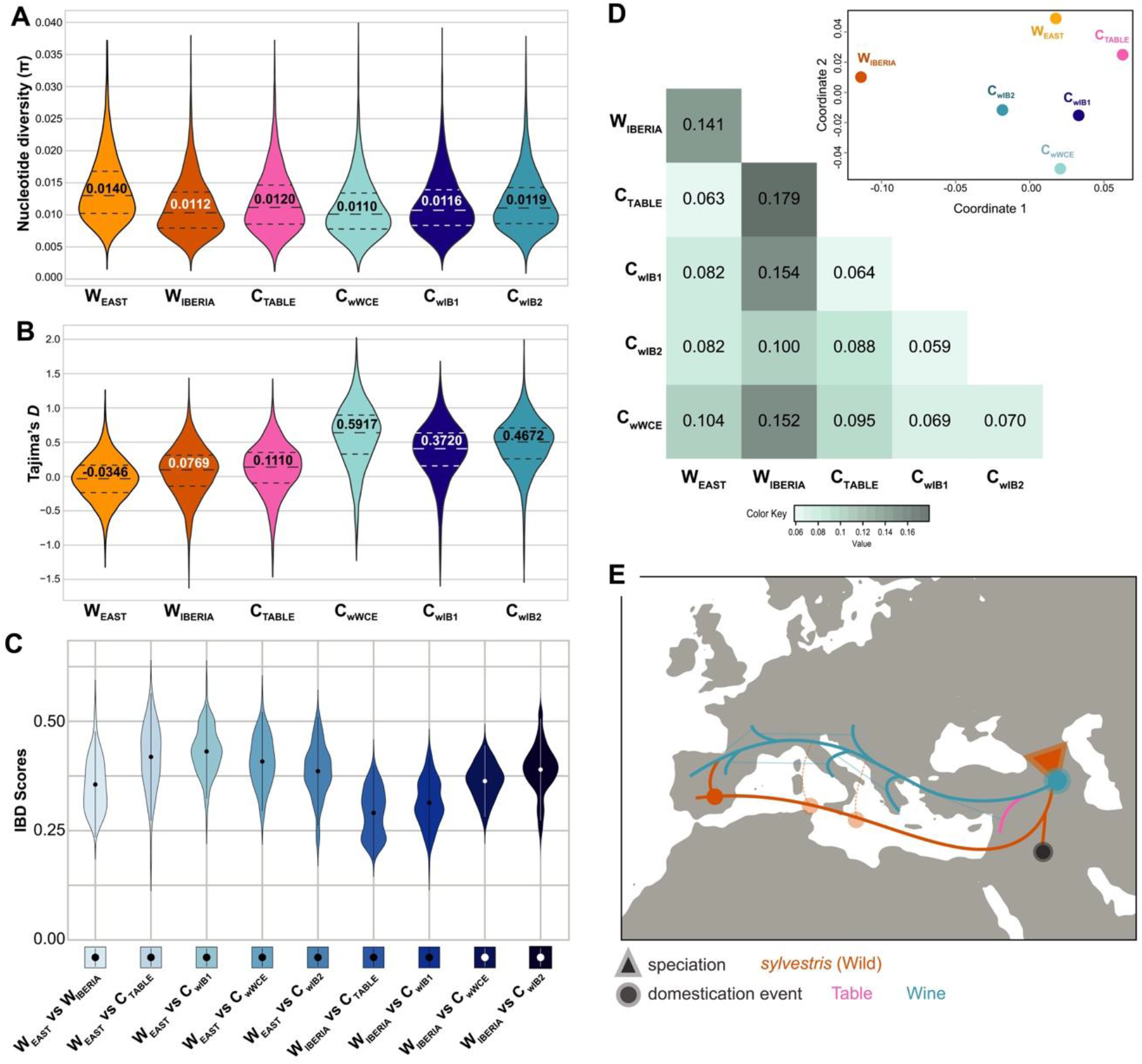
Nucleotide diversity and genetic differentiation of the six study groups. (**A,B**) Violin plot distribution of nucleotide diversity (**A**) and Tajima’s *D* (**B**). (**C**) Violin plot of pairwise IBD scores reflecting comparisons between two genotypes-of-interest. (**D**) Heat map of the group differentiation matrix of averaged F_ST_ values (*Inset*: Multidimensional scaling analysis of the F_ST_ matrix for study group differentiation). (**E**) Biogeographical model depicting *Vitis vinifera* speciation (triangle) and important events during domestication history (circles). Statistics in **A**, **B** and **D**were estimated as 100 Kb non-overlapping windows across the genome.

To capitalize on the opportunities provided by our dataset on understating grapevine domestication, we determined the impact of introgression on differentiation. We calculated the fixation index (F_*ST*_) (22) across all pairwise group comparisons and used multidimensional scaling (MDS) to reduce F_*ST*_ distances to a two-dimensional space (Fig. 3D). Results completely mirrored our previous PCA analysis (Fig. 1A). Highest F_*ST*_ was observed for table grapes against wild Iberian genotypes (F_*ST*_=0.179). This result, in conjunction with the fact that table grapes were the group closest to wild Eastern grapes (second lowest F_*ST*_=0.063), suggests an altogether independent path between table *vinifera* and Iberian *sylvestris* genotypes (Fig. 3E). Most notably, we observed three important and correlated observations: 1) all three wine groups were more differentiated against W_IBERIA_ (F_*ST*_=0.100 to 0.154) than W_EAST_ (F_*ST*_=0.082 to 0.104), suggesting a common origin in W_EAST_; 2) there was a sharp (~33%) decrease in F_*ST*_ in W_IBERIA_ vs C_wIB2_ when compared with other wine groups, supporting the existence of an introgression event; yet 3) the overall lowest F_*ST*_, with values that represent fairly low divergence, was observed between Iberian wine groups (F_ST_=0.059), as might be expected with a recent differentiation (Fig. 3D). Collectively, these observations strongly undermine the possibility of an independent grapevine domestication event based on Western European wild grape ancestors. Rather, they favor the following model: 1) a single major domestication in the Transcaucasus, 2) differentiation between wine and table grapes, 3) a post-domestication hybridization event with local wild grapes in Western Europe (most likely taking place in the Iberian Peninsula) (Fig. 3E).

### Identification of introgression tracts in Iberian grape varieties

Next, we used *f*^_*d*_ statistics to assess the fraction of the genome shared through introgression (23). The analysis was restricted to the same biogeographical space (Iberia), where we had the strongest and weakest signs of introgression in wine study groups, sympatric with the potential donor (wild) genotypes, *i.e.* configuration *f*^_*d*_ (P_1_=C_wIB1_, P_2_=C_wIB2_, P_3_=W_IBERIA_, O=*V. rotundifolia*) (Fig. 4A). Averaged genome values from the sliding-window analysis were fairly high (*f*^_*d*_=0.216). Taken together with the previous ancestry analysis (see Fig. 1C), our results suggest that the proportion of introgressed tracts may range between 25-50% of the grapevine genome in C_wIB2_ varieties. Nonetheless, we decided to implement a conservative approach in defining introgression tracts (see Methods), which resulted in 219 tracts averaging 162 Kb in size and representing 8.11% of all genomic windows (Fig. 4A). Chromosome 6 displayed the largest number of tracts (33), consistent with its highest averaged Patterson’s *D* score (Fig. 2D). In total, introgression tracts contained 2214 genes (Dataset S1). To extract biological meaning, genes were subjected to GO term functional enrichment analysis (Table S4), which highlighted an overrepresentation (*P*<0.05) in abscisic acid (ABA) signaling genes. ABA is the canonical hormone in plant adaptation to abiotic stress stimuli (24). Of significance also, is the presence of a homolog for the abscisic acid receptor *PYR1* (*VIT_02s0012g01270*), which is the sensor protein for ABA (25). Enrichment analysis also signaled grapevine *PATHOGENESIS RELATED-10* homologs, strongly associated with biotic but also abiotic stress responses (26). Results support a scenario where introgression in C_wIB2_ grapevine varieties implicated local environmental adaptation.

**Fig. 4.**
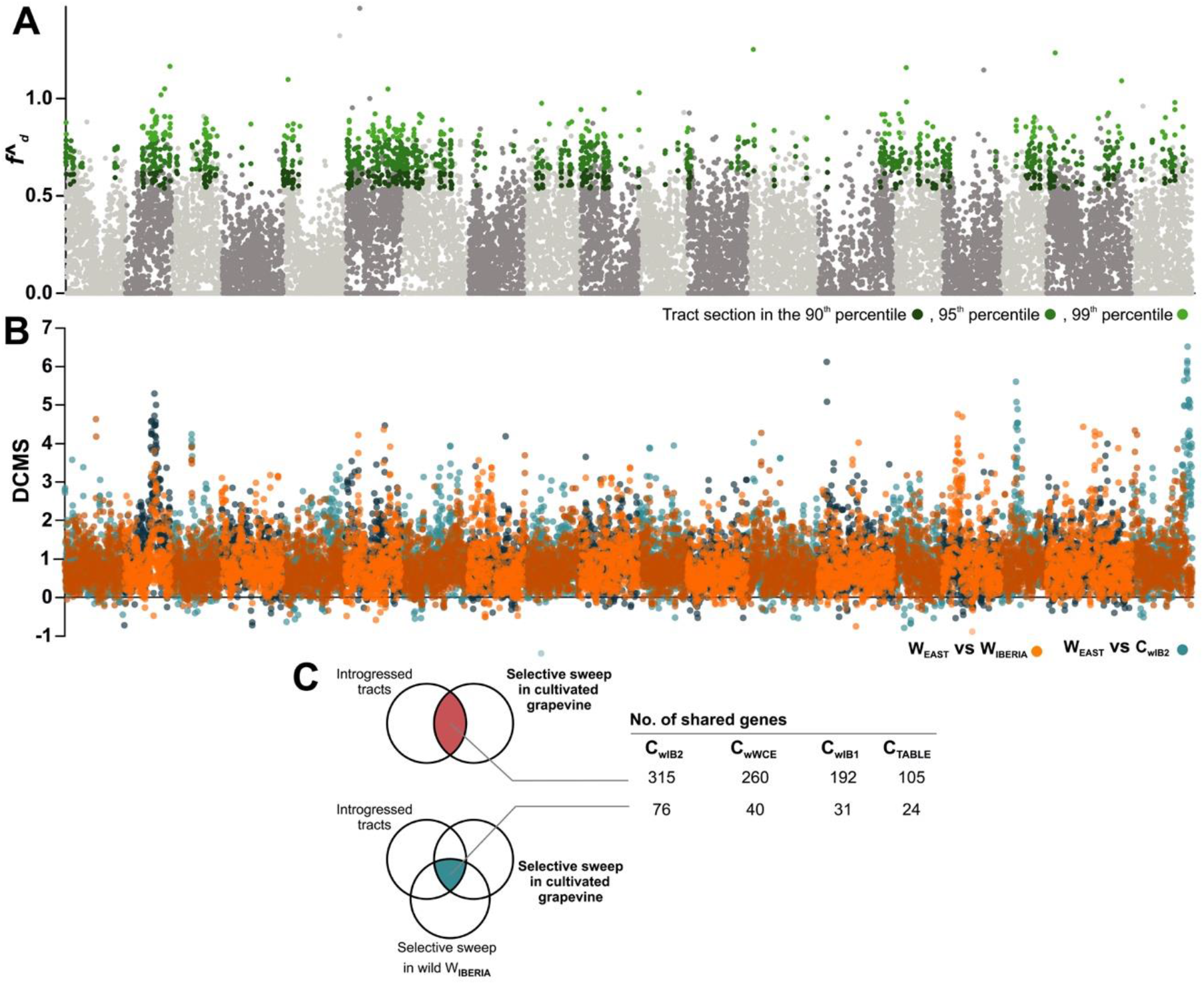
Introgression regions and signatures of positive selection in Iberian genotypes. (**A**) Manhattan plot of *f*^_*d*_ scores for detection of introgressed tracts in C_wIB2_, using 20 Kb non-overlaping windows across the genome, assuming the configuration *f*^_*d*_ (P_1_=C_wIB1_, P_2_=C_wIB2_, P_3_=W_IBERIA_, O=*V. rotundifolia*). Singleton windows with elevated *f*^_*d*_ scores were not considered as tracts (see Methods for details). (**B**) Manhattan plots of DCMS scores for W_EAST_ vs W_IBERIA_ and W_EAST_ vs C_wIB2_ comparisons, estimated across the genome in 100 Kb windows with 50 Kb steps. The *X* axis shows chromosome positions. (**C**) Venn summarization of shared genes between introgressed tracts in C_wIB2_, signatures of positive selection in wild groups (W_EAST_ vs C_wIB2_), and signatures of positive selection in cultivated grapevine groups against W_EAST_.

### Detection of signatures of positive selection across the genome and overlap with introgression tracts

We next reasoned that some regions under selection might be favored in introgressed tracts of C_wIB2_. To address this, we focused on the detection of selection signatures across the genome, by comparing cultivated varieties as well as Iberian wild genotypes, against the wild ancestral (W_EAST_) (Fig. 3E). A catalogue of potential positive selection signals was identified through four complementary statistics that utilize different properties of the data: genetic differentiation (F_*ST*_), genetic diversity (ROD), and the allele frequency spectrum of mutations (ΔTD and Fay and Wu’s H). We then implemented a summarization strategy using de-correlated composite of multiple signals (DCMS) (Table S5) (27). Multiple comparisons (Fig. 4B, Fig. S6, Dataset S2) revealed high and differentiated signals of positive selection across the genome. We observed the differentiation that underpinned domestication of wine and table grapes, with shared selection targets (*e.g.* chromosome 17) contrasting with a series of genomic regions specific of each cultivated study group (Fig. S6A-D). The data also provided an opportunity to look at the genomic fingerprint of adaptation of *sylvestris* plants as they expanded westward from their speciation center (W_EAST_ vs W_IBERIA_) (Fig. S6E). Strongly selected regions (95^th^ percentile) included an extensive set of genes associated with biotic and abiotic stress responses, including homologs of the ABA sensing and signaling pathway, and multiple pathogen Resistance genes with significance for grapevine biology (Dataset S3), indicating that biogeographical expansion was most likely driven by the capacity to adapt to newly found external challenges.

We then investigated whether introgressed tracts in C_wIB2_ might match regions under positive selection. A partial overlap could be observed between regions under selection in C_wIB2_ and W_IBERIA_ (Fig. 4B). We cross-referenced genes in introgressed tracts and genes under positive selection in Iberian wild genotypes, against multiple signals of positive selection identified for the various wine and table study groups. In this comparison, C_wIB2_ consistently displayed 2-3 times as many shared genes when compared to remaining cultivated groups (Fig. 4C). This result suggests that introgression and selection were not independent events. Furthermore, the introgressed regions contained 76 genes with signatures of positive selection in both W_IBERIA_ and C_wIB2_ (Table S6). The gene set comprised highly relevant homologs of genes associated with flowering and light perception (*CONSTANS-Like*, VIT_00s0194g00070; *FAR1-Related*, VIT_00s0194g00200), pathogen perception and hormonal signaling (*RPP2A*, VIT_18s0072g01230; *ICS2*, VIT_17s0000g05750), abiotic stress responses to cold (*ADA2B*, VIT_00s0194g00130) and drought (*ERD4*/*OSCA1.8*, VIT_02s0109g00230), and sugar content regulation (*PKR*, VIT_02s0109g00080). The latter two genes, *ERD4* and *PKR*, integrate one of the largest and most robust signals of introgression observed in our analysis, positioned in chr2, and consisting of five consecutive introgression tracts (Fig. 5). Their contiguity suggests that they may be part of a large introgression tract, containing additional positively important homologs of known abiotic and biotic stress determinants (*e.g. MED25*, VIT_02s0012g02620; *ABC-transporter Homolog*; VIT_02s0012g02770; *Disease resistance protein RPM1*, VIT_02s0012g02720; *Putative disease resistance protein RGA1*, VIT_02s0109g00420; and multiple Geraniol 8-hydroxylase-coding genes) (Fig. 5E). Most importantly, this genomic section exemplifies our hypothesis in which robust *f*^_*d*_ signals of introgression often equaled a peak in selection signatures in the C_wIB2_, including elevated levels of Tajima’s D and ROD. They also matched a peak in selection signatures in W _IBERIA_, but not in C_wIB1_ (Fig. 5C,D). *PRK* is particularly meaningful since this sugar-associated regulator is positioned within the largest introgression tract and both C_wIB2_ and W_IBERIA_ sweeps (Fig. 5E), targeting it as a key functional candidate for future characterization studies. Collectively, these findings highlight the functional pathways that most likely drove introgression from local wild populations into a range of Western European grapevine varieties.

**Fig. 5.**
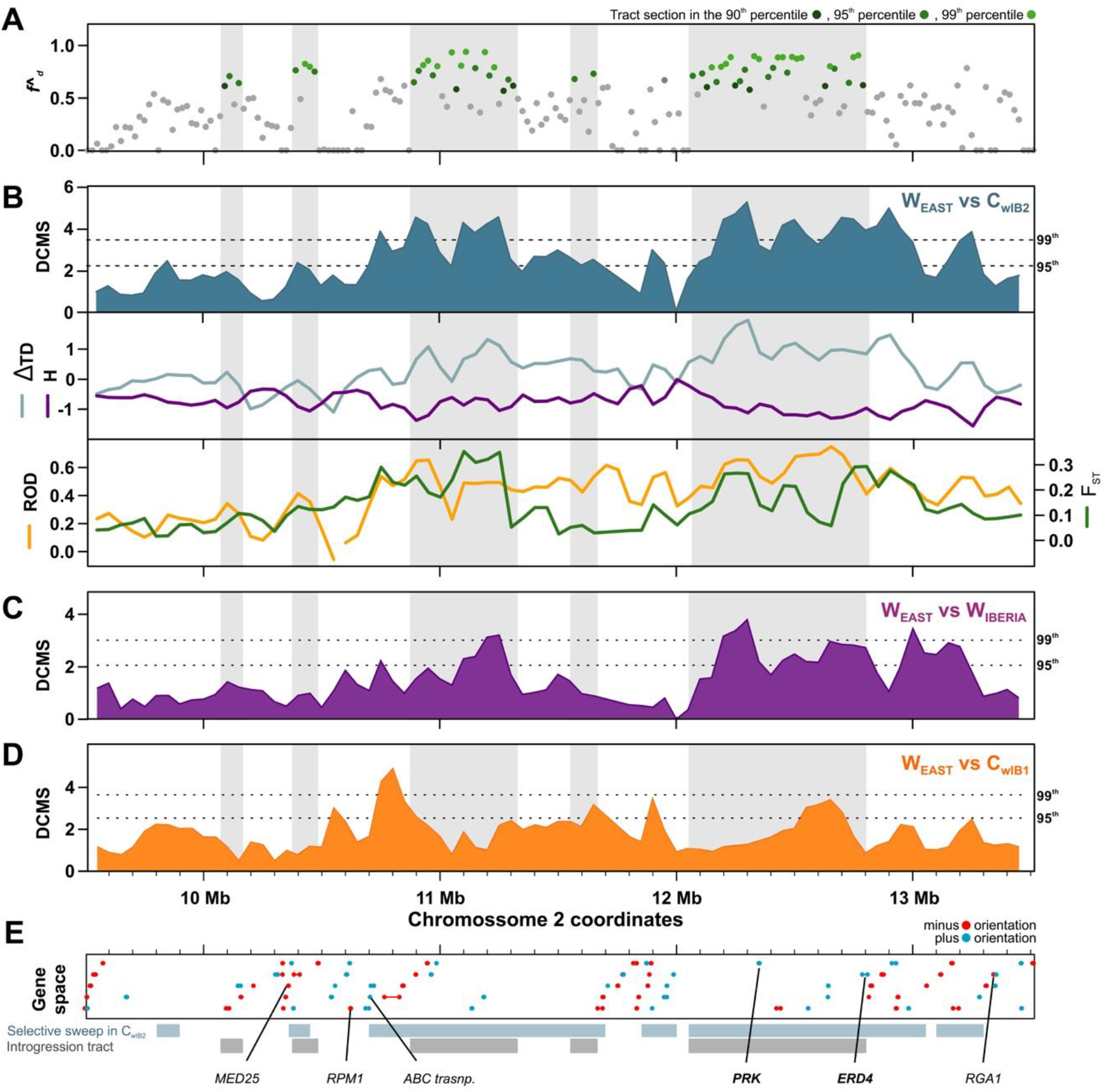
Signals of introgression and positive selection in one of the strongest introgression tracts for the C_wIB2_ study group, positioned in chromosome 2. (**A**) Zoom-in on chromosome 2 (9.5 to 13.5 Mb coordinates) details five neighboring introgression tracts determined by top *f*^_*d*_ scores in a 20 Kb sliding window analysis of configuration *f*^_*d*_ (P_1_=C_wIB1_, P_2_=C_wIB2_, P_3_=W_IBERIA_, O=*V. rotundifolia*). (**B**) DCMS scores for the W_EAST_ vs C_wIB2_ comparison (top panel), and scores for the selection signature statistics composited in the DCMS analysis (ΔTajima’s *D* and Fay and Wu’s *H* in the middle panel, F_*ST*_ and ROD in the bottom panel). (**C**) DCMS scores for the W_EAST_ vs W_IBERIA_ comparison. (**D**) DCMS scores for the W_EAST_ vs C_wIB1_ comparison. (**E**) Gene space and annotation of highlighted genes in this genomic interval.

## DISCUSSION

Implementation of whole genome resequencing is currently driving an onset of population genomic approaches that target the genetic basis of domestication, selection and adaptation events. Here, we resequenced dozens of grapevine genotypes, and provided new evidence that Western European cultivated grapes hosted a major post-domestication hybridization event with local wild genotypes, rather than independent domestication. In grapevine, such an event likely took place in the Iberian Peninsula. However, a molecular signature of introgression was also found across multiple grape varieties of Western Europe outside the Iberia. Whether a single or multiple post-domestication hybridization events occurred across the Mediterranean basin is yet to be determined. Our data supports a growing body of literature showing that hybridization events leading to post-domestication gene flow are mainstream occurrences during crop geographical expansion (28). In a few crops, this phenomenon has been associated with adaptive introgression (28, 29). Similarly, our genome-level approach allowed us to 1) recognise introgression tracts, 2) detect selective sweeps underpinning adaptation of wild populations, and 3) identify genes of interest mutually involved in adaptation and introgression.

### Western European varieties reflect post-domestication hybridization rather than independent domestication

In the present work we tested whether post-domestication hybridization or an independent domestication event underpinned the development of Western European wine varieties (reviewed by 11). The likelihood of an independent domestication event should not be minimized, since recent WGS suggests that an independent and more recent domestication event underpins a specific and nearly extinct set of grapevine cultivars from the Levant (30). However, our results provide genome-level evidence that Western European wine varieties did not originate from independent domestication, but rather from a post-domestication hybridization event. Compared to insights generated from markers with ascertainment bias, our data contradicts previous claims for an independent domestication (4, 13, 31–34), and helps clarify ambivalent hypothesis (6, 35-38), in favor of a post-domestication hybridization model (14, 39). Furthermore, our results suggest a common domestication framework that contemplates both wine and table grapes deriving from the historically and genetically accepted domestication center in the Transcaucasus, followed by expansion of grapevine across the Mediterranean basin, where it hybridized at least once with local *sylvestris* populations (Fig. 3E).

Even though feralized domesticate individuals can be found in the Iberian Peninsula, their abundance proportions seem to be fairly small (35, 37). Here, introgression directionality testing using *f*^_*d*_ statistics indicated unidirectional gene flow between *sylvestris* and *vinifera* populations, supporting previous 3-population tests on SNP chip data (14). Another key finding was the multiple evidence that all three wine study groups seem to display, to varying degrees, a *sylvestris* introgression signature (summarized as C_wIB2_>C_wWCE_>C_wIB1_) (Fig. 2). This suggests that introgression historically permeated most of the modern grapevine wine varieties found in Western Europe. Future studies should now address the origin and whether these introgression signatures derived from a single or multiple hybridization events (Fig. 3E). Our varietal dataset was structured into groups that broadly reflect previous population structuring using SNP chip data (15, 32), in which C_wIB1_ can be seen as a core group of typical Iberian varieties, while C_wIB2_ are Iberian varieties with an affinity to Western and Central Europe genotypes (15). In this context, the possibility that the hybridization responsible for C_wIB2_ took place outside the Iberian Peninsula needs to be considered. However, the likeliest scenario is that this hybridization occurred in the Iberian Peninsula. C_wIB2_ showed the highest and C_wIB1_ the lowest introgression signatures, yet both groups presented the closest genetic proximity, indicating recent differentiation. Ample studies using low resolution molecular markers support the genetic proximity between local *sylvestris* populations and modern cultivated varieties in Portugal (31, 34, 37, 40) and Spain (4, 6, 35). Finally, the chlorotype that characterizes wild populations in Western Europe is more prevalent in cultivated varieties from the Iberian Peninsula, when compared to remaining geographies (4).

In a single-introgression model, C_wIB2_-related genotypes may have acted as donors of *sylvestris* genetic material, further diluted in Western European cultivated varieties by backcrossing with *vinifera* as a result of purposeful breeding. The long track record of historical flow of varieties across Europe, particularly since Roman times (41, 42), offers a framework for interchange between Iberian and WCE varieties. Interestingly, recent ancient DNA analysis suggests a transition in French grapevine diversity from Roman to Medieval times, in which early Roman period seeds clustered closer to Iberian and Eastern European grape varieties, whereas late Roman and early Medieval seeds were more similar to modern Western Europe varieties (39). In agreement, pip morphometric studies documented a shift from abundant morphologically wild pips in earlier chronologies, to domestic types in the Late Roman and Medieval times, suggesting greater selection efforts in these later periods (42). Meanwhile, one should also consider a multiple-introgression model, in which hybridization events occurred independently in multiple geographic locations (Fig. 3E). Non-genome-level studies have suggested a genetic relatedness with local wild genotypes in cultivated varieties from Italy, France and the Balkans (6, 13, 36, 38, 43, 44), but whether these signatures reflect a single or multiple hybridization events remains to be established. Future whole genome resequencing efforts across the Mediterranean distribution range will be vital to expand our knowledge on post-domestication hybridization, particularly in the wine-producing varieties of Western Europe.

### Selection signatures corroborate a case of adaptive introgression in grapevine

Screening for wild introgression signatures that may be present in the cultivated gene pool can be an effective strategy to uncover wild diversity relevant for crop adaptation to current environmental changes (28). Our evidence suggests that adaptive introgression is common in cultivated grapes, since many wild genes under natural selection seem to have been favored and retained in introgressed cultivated varieties (Figs. 4 and 5). Here also, *D*-statistics (Fig. 2) backed previous chlorotype data (4) to support a cross with maternal *sylvestris* provenance. The size of estimated introgression in the strongest admixed study group - C_wIB2_ - suggests the replacement of a massive number of alleles for new functional variants. Such an event is likely to lead to important phenotypic differences (28). We singled out a set of 76 genes that belong to selective sweeps in C_wIB2_ as well as W_IBERIA_, and are part of introgression tracts between both groups (Table S6). They offer high confidence candidates for trait architecture determination in C_wIB2_. Using small SNP panels, Cunha and co-workers (37) recently genotyped the Portuguese national variety catalogue plus local *sylvestris* genotypes, showing that a subset of varieties with strong overlap with C_wIB2_ clustered with local wild relatives. Several varieties belonged to the Vinhos Verdes demarcated region typical of Northwestern Iberia. In our studies, 7 out of 11 of the C_wIB2_ members are canonical Vinhos Verdes varieties, firmly establishing how a major introgression event characterizes this Iberian wine type. Remarkably, Vinhos Verdes are naturally sparkling wines with lower sugar and higher acidity, suggesting that there may be a genetic component in addition to the viticultural and environmental factors that help shape their typicity. Amongst the 76 genes-of-interest (Table S6) we singled out *PKR* (VIT_02s0109g00080), due to its overlap with an interval showing strong signatures of both positive selection and introgression (Fig. 5). PKR encodes for a phosphoribulokinase that can be accounted for sugar content regulation, and its Arabidopsis best ortholog (AtPRK, AT1G32060) is involved in redox regulation of the Calvin-Benson cycle (45).

In this report we show an east-to-west genetic gradient in both wild and cultivated genotypes (PC1, Fig. 1B), which supports earlier studies (6, 14, 20, 32) and corroborates the presence of introgression. The latter is likely to have favored a reduction in the genetic load (*i.e.* the “cost of domestication”) previously reported in grapevine (9), evidenced here by the increase in nucleotide diversity in C_wIB2_. In the W_IBERIA_ population, selective sweeps were likely associated with resistance and adaptation mechanisms, detected by the presence of a large number of genes involved in transcriptional control, pathogen resistance, and hormonal modulation (Dataset S3). In other crops, adaptive introgression has been associated with adaptation to altitude and geographical expansion (46, 47). Similarly, we highlight how Vinho Verde varieties are typical of the Northwestern Iberian Peninsula, characterized by a wet Temperate Atlantic climate that contrasts severely with the dry Mediterranean climate towards the Iberia Southeast (48). Considering that *sylvestris* plants are lianas that favor high humidity conditions (35), then introgression may have enabled cultivated grapes to quickly rewire water-usage signaling and response pathways. In support, many of our 76 genes-of-interest are involved in ABA/drought responses. The Trehalose 6-Phosphate Phosphatase (TPP) family member VIT_15s0046g01000, links sugar and abiotic stress adaptation, by being involved in the control of sugar utilization as well as in the tolerance response to drought, as seen in other plants (49). VIT_18s0072g01220, is a putative grapevine ABA transporter (50) that may interfere with ABA distribution and ultimately control the ABA-regulated stress responses (51). VIT_00s0194g00210 is an ortholog of the Arabidopsis TPK1 vacuolar K+ channel involved in the ABA- and CO_2_-mediated stomatal closure (52). Other genes with orthologs implicated in drought/ABA responses include the transcriptional adapter ADA2B (VIT_00s0194g00130) (53), and Early-Responsive to Dehydration stress protein ERD4 (VIT_02s0109g00230) that belongs to the OSCA family of mechanically activated ion channels involved in osmo-sensing (54). The latter OSCA proteins were recently associated with regulation of plant stomatal immunity (55). Finally, our 76 genes-of-interest also incorporate the homologs of *ISOCHORISMATE SYNTHASE 2* (VIT_17s0000g05750), which is involved in the biosynthesis of salicylic acid, the central hormone in local and systemic acquired resistance against pathogens (56), as well as *DISEASE RESISTANCE PROTEIN RPP2A* (VIT_18s0072g01230), an ortholog of genes linked to downy mildew resistance (57). Thus, it seems that introgression impacted on upstream hormonal control of environmental responses, especially those involving ABA and SA. These results emphasize the potential of these wild populations as sources of novel allelic diversity for breeding, as suggested for grapevine and other major crops (58, 59).

### The timing of post-domestication hybridization

An important question now remains as to the historical timing of post-domestication hybridization. Genome resequencing data suggests a protracted domestication history in which *sylvestris* and *vinifera* diverged anytime between 200-400 and 22 kya. Models show an important genetic bottleneck *ca.* 8 kya that matches the earliest archaeological evidence of Eurasian grapevine use in the Transcaucasus, and marks the beginning of purposeful cultivation in grapevine (7–9, 11). It is generally accepted that the Transcaucaus region approximates the primary domestication center, after which grapevine use (and possibly a wine culture) spread to Anatolia and across the Mediterranean following the main civilizations (10, 60). This assumption provides an extensive timeframe for a subsequent hybridization event. In the Iberian Peninsula, there is evidence for gathering of wild grapes by hunter-gatherers in the Early Holocene (61), and by the first prehistoric farmers 8 to 4 kya (41). This means that locals were familiar with native *sylvestris* populations and therefore amenable to take advantage of crosses with domesticated *vinifera*. Meanwhile, the earliest evidence of grapevine cultivation in the Iberian Peninsula dates back to 2900 ya and is based on Phoenician influence (41, 62), making it the earliest tentative moment for an introgression event in this region. In stark contrast, we show how varieties from the C_wIB2_ population can display up to 25-50% of introgression tracts (Figs. 1 and 2), placing them as potential F1s or second generation backcrosses. In grapevine, this is not necessarily recent due to the extensive use of clonal lineaging since Roman times (10). In support, ancient DNA analysis of French medieval grape pips recently provided evidence for 900 y of uninterrupted vegetative propagation (39). The study also suggests the presence of gene flow between local wild grapevines and cultivated varieties, timing it to the early stages of viniculture in France (*ca.* 2500 ya). Within the context of the Iberian Peninsula, we find additional support that hybridization was not a fairly recent event. A morphometrics study of grape pips from Northwestern Iberia archaeological sites grouped medieval and Roman pips close to the modern variety *Alvarinho*(PT)/*Albariño*(SP) (63). This C_wIB2_ member is a hallmark variety for Vinhos Verdes wines, and was suggested to be a first-generation migrant from *sylvestris* based on SSR data (31). Another C_wIB2_ variety, Amaral, is mentioned in 1532 writings addressing the North of Portugal (64). Collectively, results frame a post-domestication hybridization event in the Iberian Peninsula between 2900-500 ya. Definitive clues are likely to be hidden in the DNA of archaeobotanical samples. Future approaches should concentrate on employing genomic approaches to confront ancient DNA samples with modern genomic sequences, as a means to understand the timing, strength and interdependence of hybridization events that seem to permeate a subset of Iberian varieties, and Western European varieties in general.

## MATERIALS AND METHODS

Extended Materials and Methods are available in the Supplementary Materials and Methods section.

### Sampling, sequencing and mapping

*Vinifera* varieties were sampled from two separate Portuguese germplasm collections (PORVID and UTAD), and *sylvestris* samples were collected in the southwestern region of the Iberian Peninsula. High quality genomic DNA was used to produce PCR-free sequencing libraries followed by Illumina sequencing. Sequencing data from 37 additional genotypes from previous studies were also used. Information on genotypes and sequencing effort is summarized in Table S1. Read quality filtering and trimming was performed with *FastQC* (https://www.bioinformatics.babraham.ac.uk/projects/fastqc/) and *Trimmomatic* (65). Reads were mapped to the *Vitis vinifera* PN40024 reference genome using *BWA-MEM* (66).

### Population structure analysis

For Principal Component Analysis, we estimated genotype posterior probabilities using ANGSD (16). The *ngsCovar* feature (*ngsPopGen* package) was used to compute the expected correlation matrix between individuals from genotype posterior probabilities. For the phylogenetic tree, the same genotype posterior probabilities were used to calculate pairwise genetic distances in *ngsDist* from *ngsTools*. We computed a Distance-based minimal evolution tree by inputting the genetic distance matrix into *FastME* (http://www.atgc-montpellier.fr/fastme/) with 100 bootstraps for branch support. For ancestry analysis we used *NgsAdmix* (http://www.popgen.dk/software/index.php). *NgsAdmix* was run assuming 2 to 8 ancestral populations with the default minor allele frequency of 0.05.

### Nucleotide diversity and genetic differentiation

Population genetics summary statistics (*F*_*ST*_; Watterson’s Theta, *θ*_w_; *π*; Tajima’s *D*; Fay and Wu’s *H*) were also inferred under a probabilistic framework using ANGSD. *V. rotundifolia* (9) was used as the outgroup to polarize the ancestral state of alleles at each polymorphic site. All statistics were summarized across the genome using a sliding-window approach.

### IBD estimation and SNP calling

To look at the relationship between different grape cultivars, we performed Identity-By-Descent (IBD) analysis using the probabilistic methods implemented in ANGSD (16). We applied stringent criteria for the SNP call, with post-cutoff of 0.95 and a SNP *P* value of 1×10^−9^, to include only highly supported SNPs. Subsequently, the SNPs were used to calculate IBD for all pairwise comparisons among the 100 samples using PLINK (67) and applying the following filters: maf 0.05 and geno 0.05.

### Admixture test using Patterson’s *D* statistic

Patterson’s *D* statistics (or ABBA-BABA test) assumes three populations (P_1_,P_2_,P_3_) and one outgroup (O), which are phylogenetically related as (((P_1_,P_2_),P_3_),O). Here, we estimated all permutations of the six groups of interest (W_EAST_, W_IBERIA_, C_TABLE_, C_wWCE_, C_wIB1_, C_wIB2_) as P_1_, P_2_ and P_3_. *Vitis rotundifolia* served as an outgroup (O). We computed Patterson’s *D* statistic using allele frequencies instead of binary counts of fixed ABBA-BABA sites, as implemented in the *ABBABABA2 (Multipopulation)* function in ANGSD (16), using non-overlapping 20 Kbp windows. We calculated *f*^_*d*_ to assess the fraction of the genome shared through introgression (23) in the comparison P_1_=C_wIB1_, P_2_=C_wIB2_, P_3_=W_IBERIA_, O=*V. rotundifolia*. We estimated *D*(P_1_,P_2_,P_2_,O) and *D*(P_1_,P_3_,P_3_,O) in ANGSD as previously described, which allowed us to obtain, for each genomic window, a donor population P_D_ (population with the higher frequency of the derived allele).

### DCMS analysis of positive selection signatures

We calculated four separate statistics that differ in their approach to detect selection events based on the type of selection signals that are targeted: genetic differentiation (F_*ST*_), shifts in the allele frequency spectrum of mutations (Delta Tajimas’s *D* - ΔT*D* - and Fay and Wu’s *H*) and reduction of genetic diversity from pairwise nucleotide diversity measures (reduction of diversity - ROD). Statistics were based on F_*ST*_, Tajima’s *D*, Fay and Wu’s *H*, and nucleotide diversity (*π*) calculated across the genome (100 Kbp windows, 50 kb steps) as previously reported. De-correlated composite of multiple signals (DCMS) was then used to summarize the four different statistics, taking the covariance of the statistics into account (27). Comparisons-of-interest were defined as those that confronted the *W_EAST_* group with the remaining five groups. Genes-of-interest were considered for genomic windows above the 95^th^ percentile of the distribution.

### Analysis of genes of interest

Gene annotation was retrieved from PANTHER (http://pantherdb.org) and UniProtKB (https://www.uniprot.org/uniprot/). Also, for the 76 crossreferenced genes-of-interest, protein Fasta sequences were retrieved from UniProtKB, and used to perform a BlastP search in NCBI (https://blast.ncbi.nlm.nih.gov/Blast.cgi), against the *Non-redundant protein sequences (nr)* database, and the *Arabidopsis Thaliana RefSeq* database. GO terms (*GO biological process complete*) were subjected to statistical overrepresentation testing in PANTHER (http://www.pantherdb.org).

## ACKNOWLEDGMENTS

This work was funded by the Norte Portugal Regional Operational Programme (NORTE 2020), through the European Regional Development Fund (FEDER) [NORTE-01-0145-FEDER-000007]. Financial support was provided by Fundação para a Ciência e Tecnologia (FCT/MCTES) to S.F. [SFRH/BD/120020/2016], M.A.G. [PD/BD/114042/2015], A.J.M.P. [SFRH/BPD/111015/2015], I.C. [UIDB/04033/2020], P.H.C. [PTDC/BAA-AGR/31122/2017], M.C. [CEECINST/00014/2018] and H.A. [CEECIND/00399/2017].

## SUPPLEMENTARY FIGURES

**Fig. S1.**
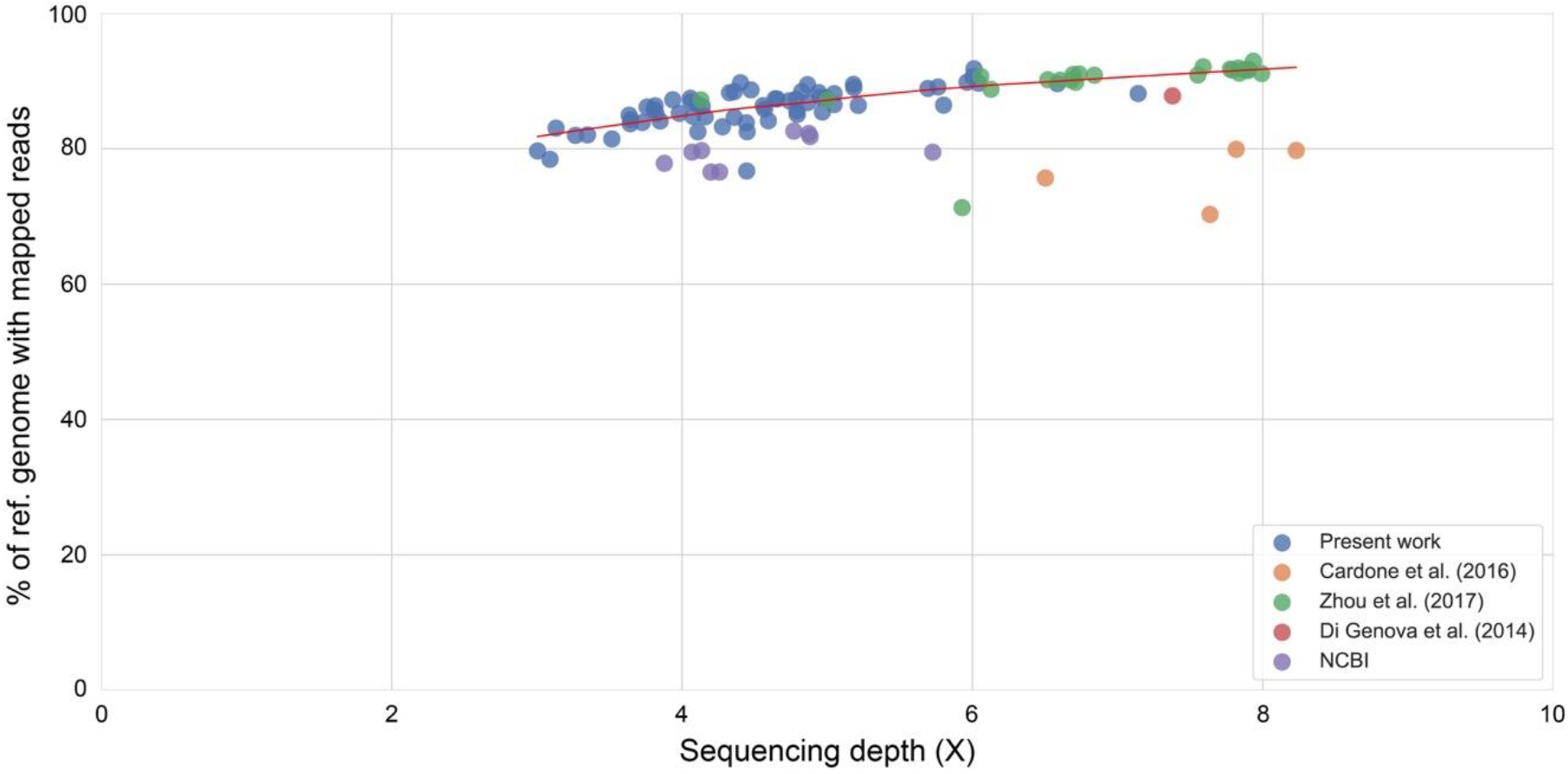
Percentage of the reference genome with at least one mapped read, plotted as a function of the sample’s sequencing depth.

**Fig. S2.**
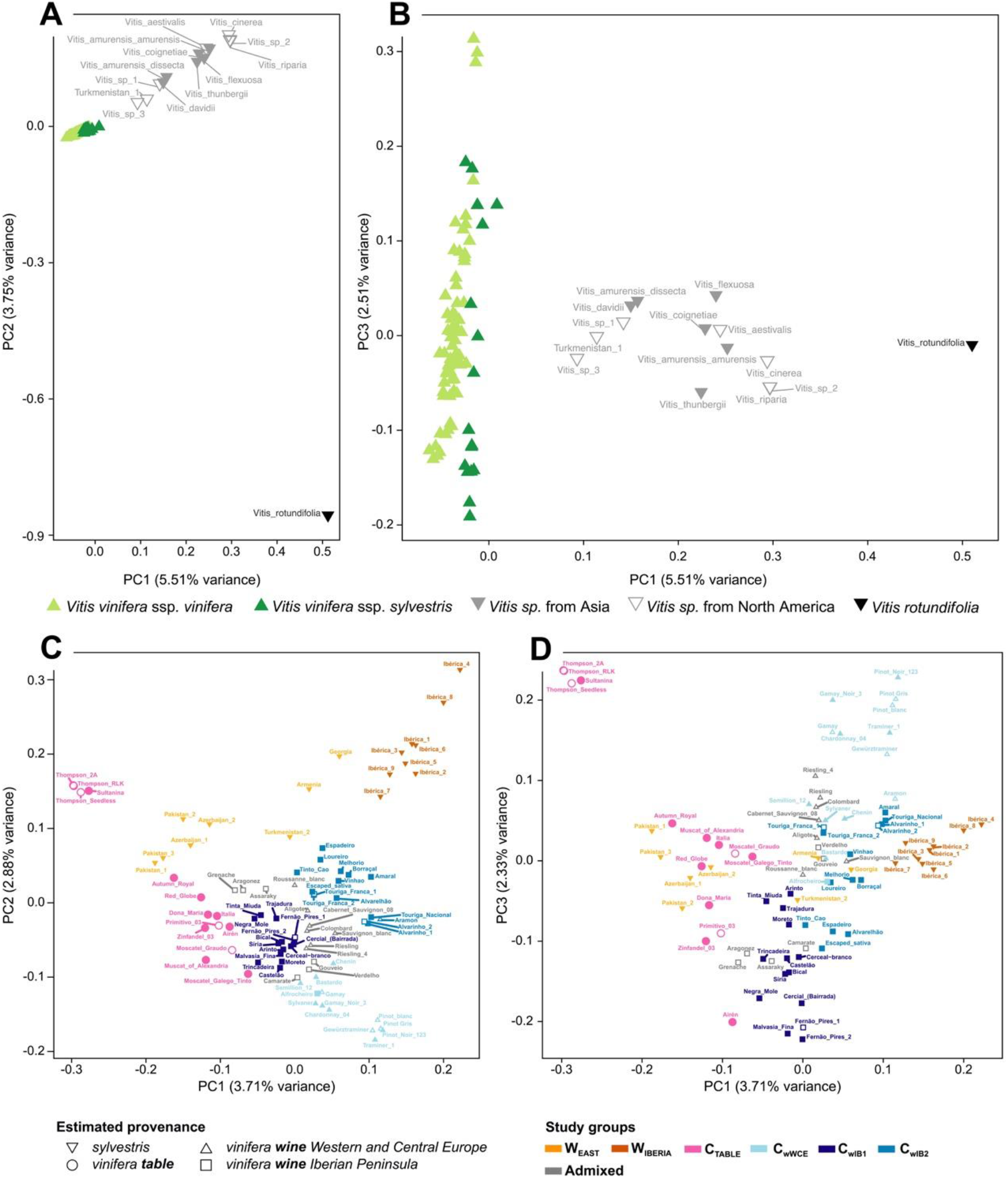
Detailed PCA plots for population structure analysis. PCA of the 100 sampled genotypes [*Vitis* sp. as well as wild (*sylvestris*) and cultivated (*vinifera*) grapevine genotypes], for eigenvectors 1 and 2 (**A**) and 1 and 3 (**B**). PCA of *Vitis vinifera* wild and cultivated genotypes, for eigenvectors 1 and 2 (**C**) and 1 and 3 (**D**). In **C,D**, open symbols depict genotypes excluded from the six study groups due to admixture or clonal redundancy.

**Fig. S3.**
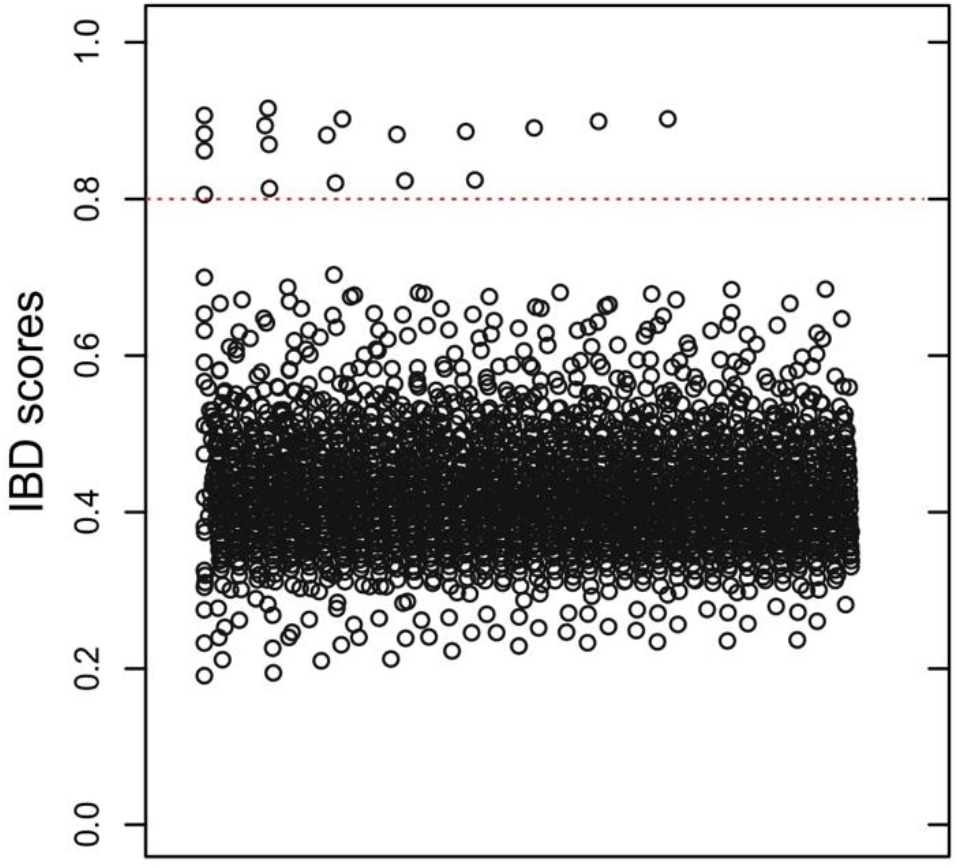
Identical-by-Descent (IBD) pairwise analysis of cultivated grapevine varieties. In total, 23 genotypes evidenced clonal relationships (Table S2). Robustness of the analysis was validated by the identification, within clonal pairs, of: 1) three clone pairs (Fernão_Pires, Touriga_Franca and Alvarinho) purposefully incorporated into the sampling effort; 2) well established sports (e.g. Pinot Noir, Pinot Gris, Pinot Blanc). Since presence of clones will skew allele frequencies specific to an individual, ultimately altering study group and nucleotide diversity statistics, just one of the clones was incorporated into the six study groups defined in the present study. Dashed line represented the threshold for assignment of clonal nature between pairwise compared genotypes. All 18 comparisons with IBD>0.8 represented expected clones.

**Fig. S4.**
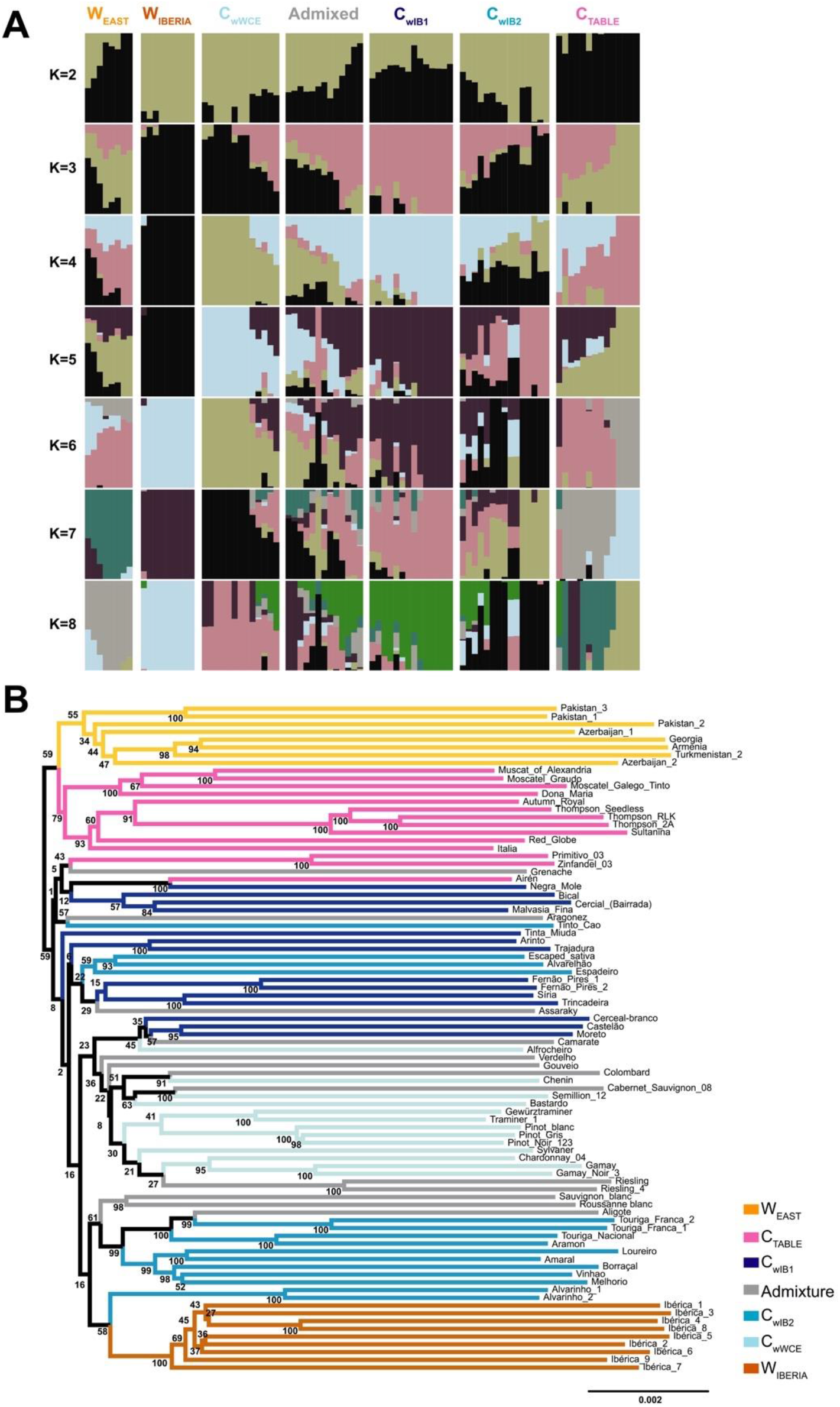
Ancestry and phylogenetic tree analysis of population structure. (**A**) Ancestry proportions of all *Vitis vinifera* genotypes following admixture analysis for K equaling to 2–8. (**B**) Phylogenetic tree of *Vitis vinifera* genotypes, with bootstrap values indicated for all nodes (100 replicates).

**Fig. S5.**
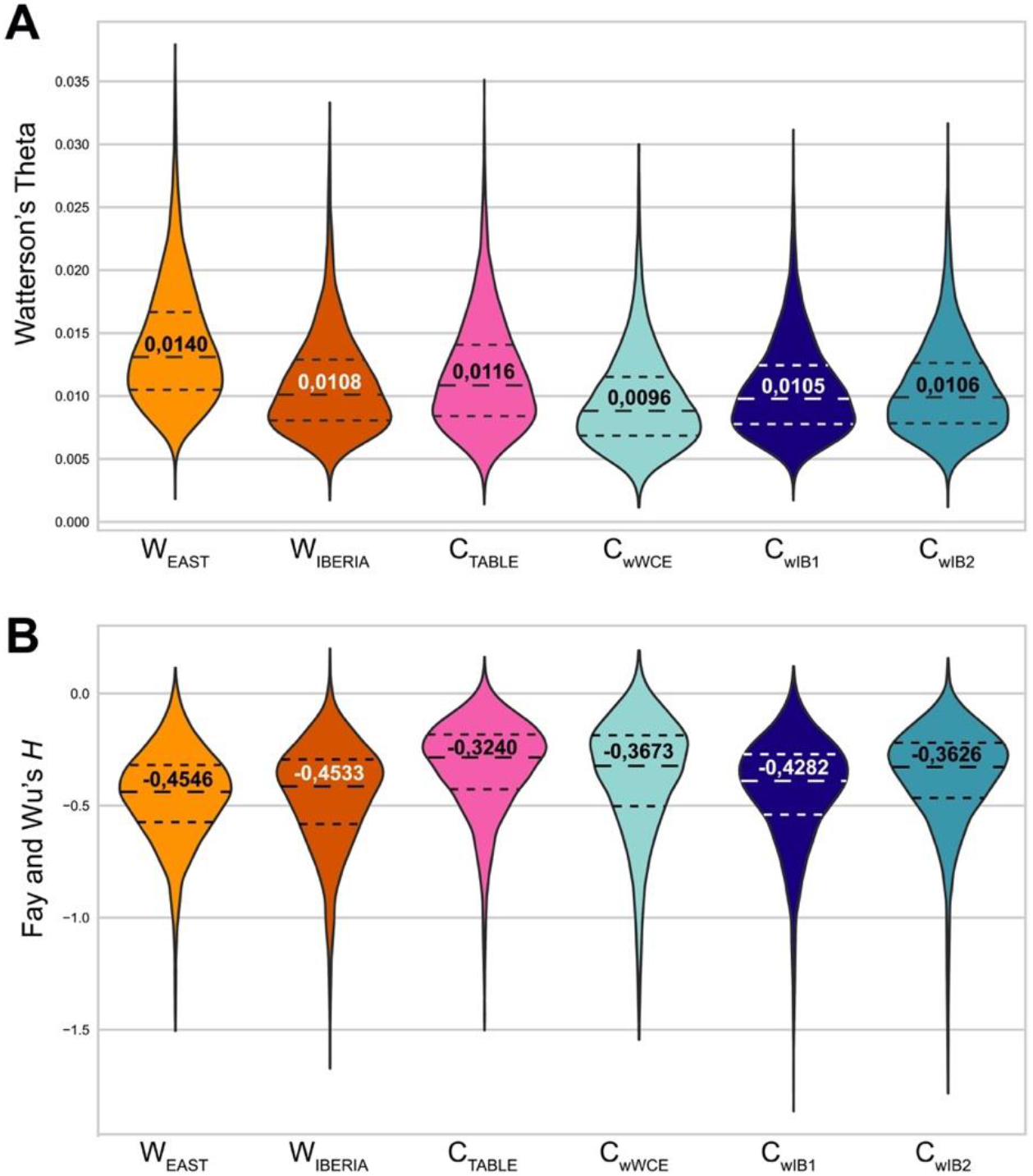
Nucleotide diversity and genetic differentiation of the six study groups. Violin plot distribution of Watterson’s Theta (**A**) and Fay and Hu’s *H* (**B**).

**Fig. S6.**
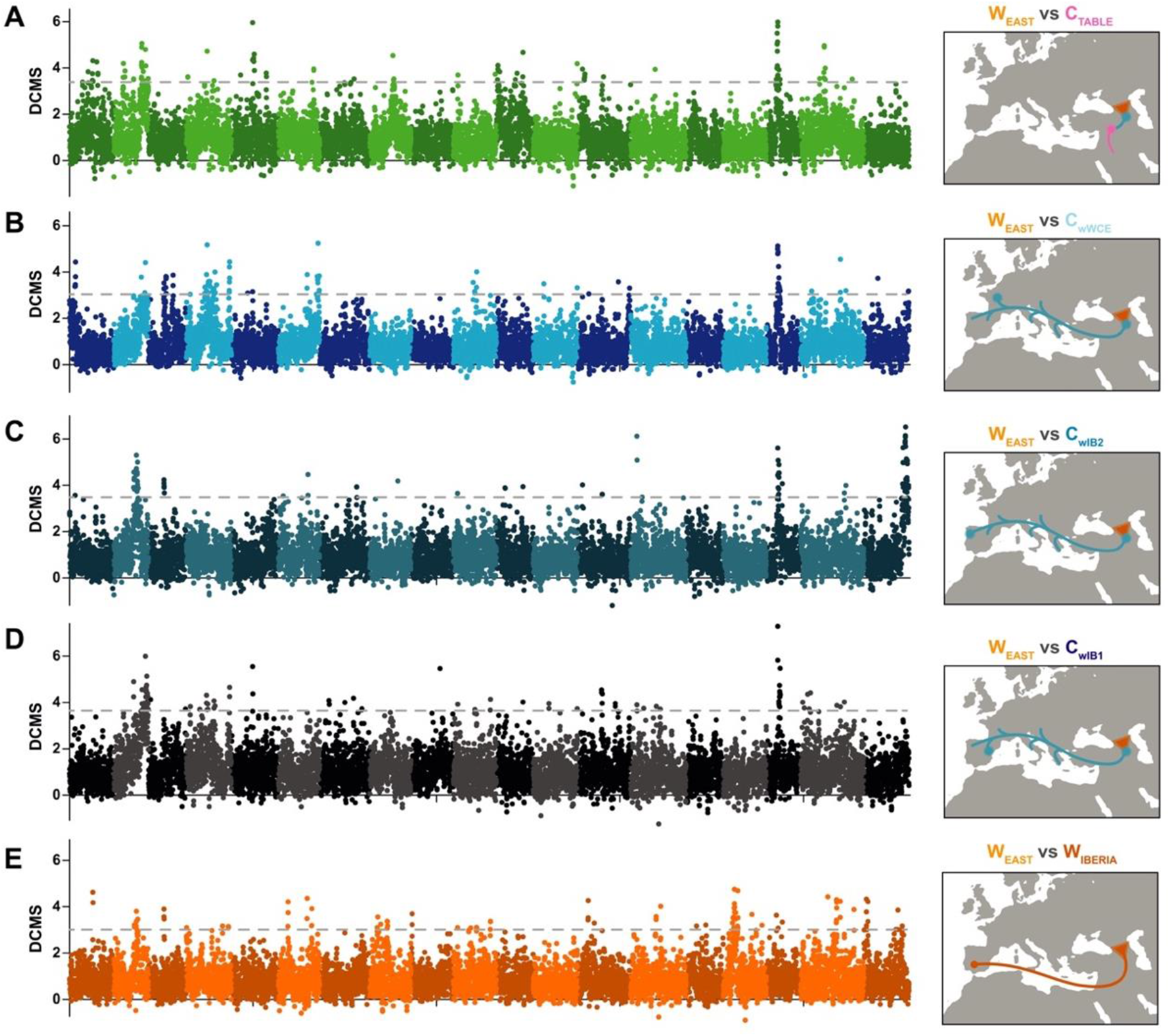
Signatures of positive selection for wild and cultivated study groups when compared against W_EAST_. Manhattan plots of DCMS scores for C_TABLE_ vs W_EAST_ (**A**), C_wWCE_ vs W_EAST_ (**B**), C_wIB2_ vs W_EAST_ (**C**), C_wIB1_ vs W_EAST_ (**D**) and W_IBERIA_ vs W_EAST_ (**E**), estimated across the genome in 100 Kbp windows with 50 Kbp steps. Dashed line represents 95^th^ percentile cut-off. The *X* axis shows chromosome positions.

## SUPPLEMENTARY TABLES

**Table S1.**
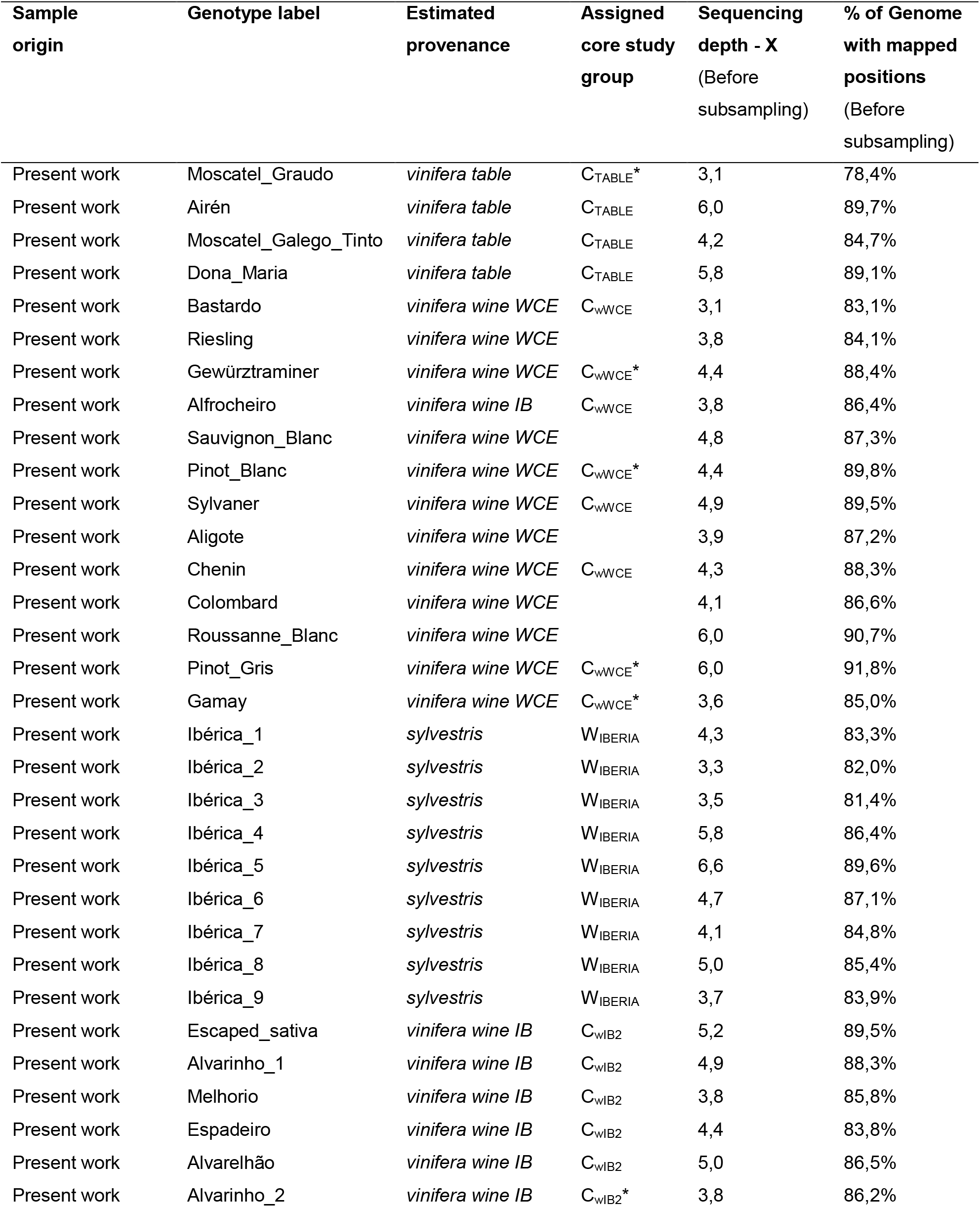
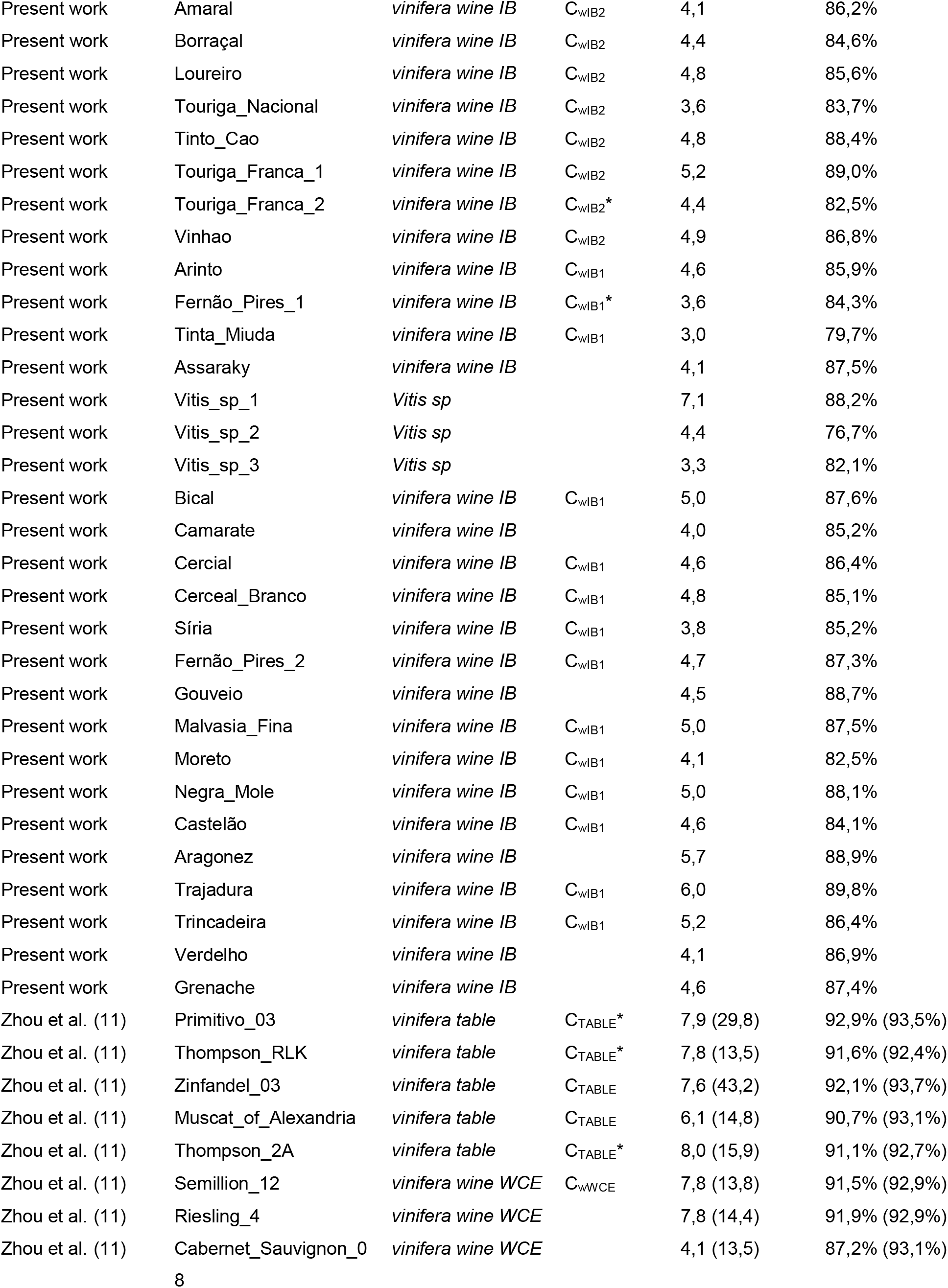
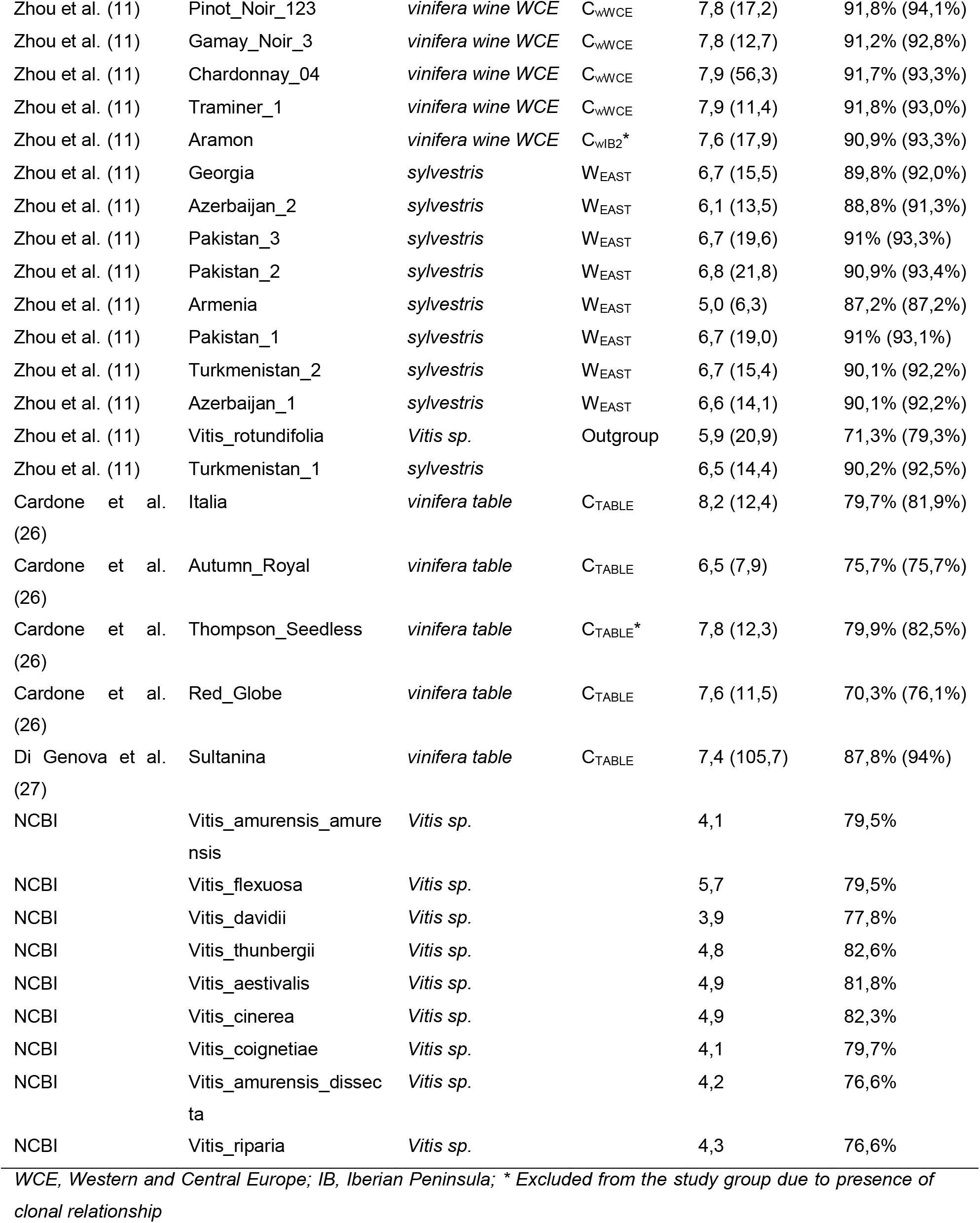
Plant material and sequencing summary statistics.

**Table S2.** List of clonal relationships between genotypes, identified by IBD analysis.

| Clones | Purposefully incorporated<br>clonal pairs |
| --- | --- |
| Sultanina, Thompson_Seedless; Thompson_2A; Thompson_RLK |  |
| Pinot_Noir_123; Pinot_Gris; Pinot_Blanc |  |
| Zinfandel_03; Primitivo_03 |  |
| Touriga_Nacional, Aramon |  |
| Fernão_Pires_2; Fernão_Pires_1 | × |
| Muscat_of_Alexandria; Moscatel_Graudo |  |
| Traminer_1; Gewürztraminer |  |
| Alvarinho_1; Alvarinho_2 | × |
| Touriga_Franca_1; Touriga_Franca_2 | × |
| Riesling_4; Riesling |  |
| Gamay_Noir3; Gamay |  |

**Table S3.** Genome-wide results and significance of selected comparisons for Patterson’s *D* statistic.

| **P<sub>1</sub>** | **P<sub>2</sub>** | **P<sub>3</sub>** | ***D* Statistic** | **Z score** | ***P* value** |
| --- | --- | --- | --- | --- | --- |
| C<sub>TABLE</sub> | C<sub>wIB2</sub> | W<sub>EAST</sub> | -0.033087 | -25.539736 | 0.000000 |
| C<sub>TABLE</sub> | C<sub>wIB2</sub> | W<sub>IBERIA</sub> | 0.177209 | 114.249097 | 0.000000 |
| C<sub>TABLE</sub> | C<sub>wWCE</sub> | W<sub>EAST</sub> | -0.026247 | -17.966964 | 0.000000 |
| C<sub>TABLE</sub> | C<sub>wWCE</sub> | W<sub>IBERIA</sub> | 0.144810 | 76.765866 | 0.000000 |
| C<sub>TABLE</sub> | C<sub>wIB1</sub> | W<sub>EAST</sub> | -0.011705 | -9.555438 | 0.000000 |
| C<sub>TABLE</sub> | C<sub>wIB1</sub> | W<sub>IBERIA</sub> | 0.092798 | 59.475533 | 0.000000 |
| C<sub>wIB1</sub> | C<sub>wIB2</sub> | W<sub>EAST</sub> | -0.023259 | -20.448190 | 0.000000 |
| C<sub>wIB1</sub> | C<sub>wIB2</sub> | W<sub>IBERIA</sub> | 0.094978 | 69.026545 | 0.000000 |
| C<sub>wIB1</sub> | C<sub>wWCE</sub> | W<sub>EAST</sub> | -0.015487 | -11.816794 | 0.000000 |
| C<sub>wIB1</sub> | C<sub>wWCE</sub> | W<sub>IBERIA</sub> | 0.056776 | 33.590551 | 0.000000 |
| C<sub>wIB2</sub> | C<sub>wWCE</sub> | W<sub>EAST</sub> | 0.008206 | 6.243488 | 0.000000 |
| C<sub>wIB2</sub> | C<sub>wWCE</sub> | W<sub>IBERIA</sub> | -0.040772 | -25.183210 | 0.000000 |

**Table S4.** GO term statistical enrichment for Biological Process, of genes present within introgression tracts of the *C*_*wIB2*_ study group.

| GO Biological process category;<br>GO term hierarchy is represented | Observed<br>no. of<br>genes | Expected<br>no. of<br>genes | Fold<br>enrichment | FDR<br>corrected<br>*P* value |
| --- | --- | --- | --- | --- |
| **abscisic acid-activated signaling pathway (GO:0009738)** | **18** | **3.31** | **5.43** | **2.19E-04** |
| cellular response to abscisic acid stimulus (GO:0071215) | 19 | 3.75 | 5.06 | 1.83E-04 |
| cellular response to alcohol (GO:0097306) | 19 | 3.75 | 5.06 | 1.46E-04 |
| response to alcohol (GO:0097305) | 28 | 9.72 | 2.88 | 1.18E-03 |
| response to oxygen-containing compound (GO:1901700) | 49 | 27.61 | 1.78 | 4.80E-02 |
| cellular response to organic substance (GO:0071310) | 42 | 21.50 | 1.95 | 2.89E-02 |
| cellular response to oxygen-containing compound (GO:1901701) | 33 | 11.63 | 2.84 | 3.03E-04 |
| cellular response to hormone stimulus (GO:0032870) | 37 | 15.46 | 2.39 | 2.27E-03 |
| cellular response to endogenous stimulus (GO:0071495) | 37 | 16.56 | 2.23 | 6.93E-03 |
| cellular response to lipid (GO:0071396) | 25 | 6.11 | 4.09 | 1.38E-04 |
| response to lipid (GO:0033993) | 33 | 12.96 | 2.55 | 1.58E-03 |
| response to abscisic acid (GO:0009737) | 28 | 9.72 | 2.88 | 1.12E-03 |
| hormone-mediated signaling pathway (GO:0009755) | 36 | 14.87 | 2.42 | 1.69E-03 |
| **regulation of protein serine/threonine phosphatase activity (GO:0080163)** | **17** | **2.28** | **7.45** | **4.03E-05** |
| regulation of phosphoprotein phosphatase activity (GO:0043666) | 21 | 5.67 | 3.70 | 7.89E-04 |
| regulation of protein dephosphorylation (GO:0035304) | 21 | 5.67 | 3.70 | 8.46E-04 |
| regulation of dephosphorylation (GO:0035303) | 21 | 5.96 | 3.52 | 1.22E-03 |
| regulation of phosphate metabolic process (GO:0019220) | 31 | 14.21 | 2.18 | 2.92E-02 |
| regulation of phosphorus metabolic process (GO:0051174) | 31 | 14.28 | 2.17 | 4.23E-02 |
| regulation of protein modification process (GO:0031399) | 33 | 15.75 | 2.09 | 4.89E-02 |
| regulation of phosphatase activity (GO:0010921) | 21 | 5.82 | 3.61 | 1.04E-03 |
| **negative regulation of phosphoprotein phosphatase activity (GO:0032515)** | **17** | **2.94** | **5.77** | **1.05E-04** |
| negative regulation of protein dephosphorylation<br>(GO:0035308) | 17 | 2.94 | 5.77 | 1.22E-04 |
| negative regulation of protein modification process<br>(GO:0031400) | 20 | 4.34 | 4.60 | 1.21E-04 |
| negative regulation of cellular protein metabolic<br>process (GO:0032269) | 22 | 8.91 | 2.47 | 4.92E-02 |
| negative regulation of protein metabolic process<br>(GO:0051248) | 22 | 8.91 | 2.47 | 4.77E-02 |
| negative regulation of dephosphorylation (GO:0035305) | 17 | 3.02 | 5.63 | 1.08E-04 |
| negative regulation of phosphate metabolic process<br>(GO:0045936) | 19 | 4.12 | 4.61 | 1.98E-04 |
| negative regulation of phosphorus metabolic<br>process (GO:0010563) | 19 | 4.12 | 4.61 | 2.16E-04 |
| negative regulation of phosphatase activity (GO:0010923) | 17 | 3.02 | 5.63 | 1.21E-04 |
| negative regulation of hydrolase activity (GO:0051346) | 18 | 5.23 | 3.44 | 6.05E-03 |
| negative regulation of catalytic activity (GO:0043086) | 27 | 11.04 | 2.45 | 2.29E-02 |
| negative regulation of molecular function<br>(GO:0044092) | 27 | 11.26 | 2.40 | 2.51E-02 |
| **lipid biosynthetic process (GO:0008610)** | **59** | **32.54** | **1.81** | **9.94E-03** |

**Table S5.**
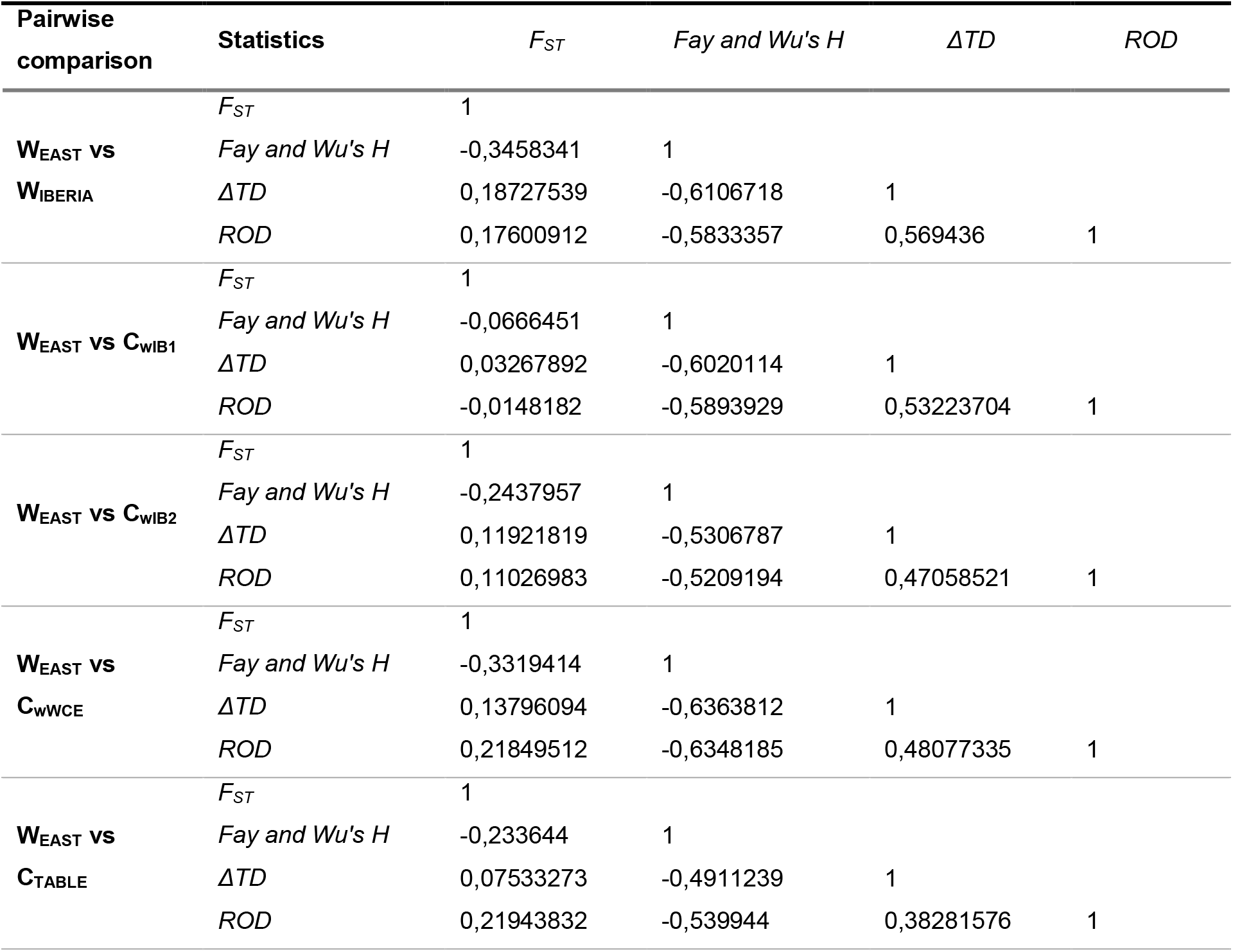
Correlation values estimated for the four statistics employed in DCMS analysis of two comparisons of interest.

| Pairwise comparison | Statistics | $F_{ST}$ | Fay and Wu's $H$ | $\Delta TD$ | ROD |
| --- | --- | --- | --- | --- | --- |
| **W<sub>EAST</sub> vs W<sub>IBERIA</sub>** | $F_{ST}$ | 1 | | | |
| | Fay and Wu's $H$ | -0,3458341 | 1 | | |
| | $\Delta TD$ | 0,18727539 | -0,6106718 | 1 | |
|  | ROD | 0,17600912 | -0,5833357 | 0,569436 | 1 |
| **W<sub>EAST</sub> vs C<sub>WIB1</sub>** | $F_{ST}$ | 1 | | | |
| | Fay and Wu's $H$ | -0,0666451 | 1 | | |
| | $\Delta TD$ | 0,03267892 | -0,6020114 | 1 | |
|  | ROD | -0,0148182 | -0,5893929 | 0,53223704 | 1 |
| **W<sub>EAST</sub> vs C<sub>WIB2</sub>** | $F_{ST}$ | 1 | | | |
| | Fay and Wu's $H$ | -0,2437957 | 1 | | |
| | $\Delta TD$ | 0,11921819 | -0,5306787 | 1 | |
|  | ROD | 0,11026983 | -0,5209194 | 0,47058521 | 1 |
| **W<sub>EAST</sub> vs C<sub>WWCE</sub>** | $F_{ST}$ | 1 | | | |
| | Fay and Wu's $H$ | -0,3319414 | 1 | | |
| | $\Delta TD$ | 0,13796094 | -0,6363812 | 1 | |
|  | ROD | 0,21849512 | -0,6348185 | 0,48077335 | 1 |
| **W<sub>EAST</sub> vs C<sub>TABLE</sub>** | $F_{ST}$ | 1 | | | |
| | Fay and Wu's $H$ | -0,233644 | 1 | | |
| | $\Delta TD$ | 0,07533273 | -0,4911239 | 1 | |
|  | ROD | 0,21943832 | -0,539944 | 0,38281576 | 1 |

**Table S6.** Annotation of 76 genes shared between the genomic regions showing simultaneous signs of introgression, positive selection in W_IBERIA_ and positive selection in C_wIB2_.

| Gene ID | Annotation | Chr | Start | End |
| --- | --- | --- | --- | --- |
| VIT_00s0194g00070 | ZINC FINGER PROTEIN CONSTANS-LIKE 9 | 10 | 2260252 | 2272206 |
| VIT_00s0194g00080 | PHOTOTROPIC-RESPONSIVE NPH3 FAMILY PROTEIN | 10 | 2272444 | 2275038 |
| VIT_00s0194g00090 | KETOACYL-ACP SYNTHASE 1 | 10 | 2280897 | 2285357 |
| VIT_00s0194g00100 | C2 AND GRAM DOMAIN-CONTAINING PROTEIN | 10 | 2288893 | 2297485 |
| VIT_00s0194g00110 | 2-PHOSPHOGLYCOLATE PHOSPHATASE 2 | 10 | 2300759 | 2305096 |
| VIT_00s0194g00120 | SERINE ACETYLTRANSFERASE 1, CHLOROPLASTIC | 10 | 2305715 | 2307059 |
| VIT_00s0194g00130 | TRANSCRIPTIONAL ADAPTER ADA2B | 10 | 2313677 | 2318858 |
| VIT_00s0194g00140 | TRAPPC3 | 10 | 2319562 | 2321948 |
| VIT_00s0194g00150 | TRAPPC3 | 10 | 2323170 | 2323607 |
| VIT_00s0194g00160 | CYTOCHROME P450, FAMILY 707, SUBFAMILY A, POLYPEPTIDE 1 | 10 | 2341847 | 2342704 |
| VIT_00s0194g00170 | KILLING ME SLOWLY 2 | 10 | 2345752 | 2346605 |
| VIT_00s0194g00180 | HYPOTHETICAL PROTEIN | 10 | 2351931 | 2352567 |
| VIT_00s0194g00190 | PUTATIVE ENDONUCLEASE OR GLYCOSYL HYDROLASE | 10 | 2352998 | 2353916 |
| VIT_00s0194g00200 | FAR1-RELATED SEQUENCE 10 ISOFORM 1 | 10 | 2358663 | 2360918 |
| VIT_00s0194g00210 | OUTWARD RECTIFYING POTASSIUM CHANNEL PROTEIN | 10 | 2364396 | 2365775 |
| VIT_00s0194g00220 | PROTEASOME ACTIVATING PROTEIN 200 | 10 | 2366617 | 2367043 |
| VIT_00s0194g00230 | PROTEASOME SUBUNIT PAB1 | 10 | 2369713 | 2370453 |
| VIT_00s0194g00240 | UB-LIKE PROTEASE 1A | 10 | 2395023 | 2395911 |
| VIT_02s0012g02920 | ACYL-COA OXIDASE 6 | 2 | 11133984 | 11136347 |
| VIT_02s0012g02970 | UNCHARACTERIZED PROTEIN | 2 | 11183822 | 11186832 |
| VIT_02s0109g00080 | PHOSPHORIBULOKINASE | 2 | 12348163 | 12351903 |
| VIT_02s0109g00130 | UNCHARACTERIZED PROTEIN | 2 | 12422839 | 12441134 |
| VIT_02s0109g00160 | UNCHARACTERIZED PROTEIN | 2 | 12557348 | 12557838 |
| VIT_02s0109g00180 | AMINOPHOSPHOLIPID ATPASE 2 | 2 | 12640920 | 12641306 |
| VIT_02s0109g00190 | ABC TRANSPORTER-LIKE PROTEIN | 2 | 12642481 | 12643852 |
| VIT_02s0109g00230 | EARLY-RESPONSIVE TO DEHYDRATION STRESS PROTEIN (ERD4) | 2 | 12786746 | 12808921 |
| VIT_06s0004g04370 | HISTONE H4 | 6 | 5352023 | 5352367 |
| VIT_06s0004g04380 | MITOCHONDRIAL F1F0-ATP SYNTHASE | 6 | 5357240 | 5358176 |
| VIT_06s0004g04390 | UDP-XYL SYNTHASE 6 | 6 | 5358853 | 5364485 |
| VIT_06s0004g04400 | P-HYDROXYBENZOIC ACID EFFLUX PUMP SUBUNIT | 6 | 5365814 | 5368974 |
| VIT_06s0004g04410 | SEC14 CYTOSOLIC FACTOR FAMILY<br>PROTEIN / PHOSPHOGLYCERIDE TRANSFER<br>FAMILY PROTEIN | 6 | 5375905 | 5383629 |
| VIT_06s0004g04420 | METALLOTHIOL TRANSFERASE FOSB | 6 | 5386184 | 5387156 |
| VIT_06s0004g04430 | UBIQUITIN-PROTEIN LIGASE 7 | 6 | 5387285 | 5399913 |
| VIT_06s0004g04440 | PATHOGENESIS-RELATED THAUMATIN<br>SUPERFAMILY PROTEIN | 6 | 5401080 | 5402253 |
| VIT_06s0004g04450 | PHOSPHATIDYLINOSITOL-SPECIFIC<br>PHOSPHOLIPASE C4 | 6 | 5406399 | 5406925 |
| VIT_06s0004g04460 | INNER MEMBRANE PROTEIN ALBINO3,<br>CHLOROPLASTIC | 6 | 5407544 | 5415216 |
| VIT_06s0004g04470 | HEAT SHOCK COGNATE PROTEIN 70 | 6 | 5418191 | 5420677 |
| VIT_06s0004g04480 | HSP20-LIKE CHAPERONES SUPERFAMILY<br>PROTEIN ISOFORM 1 | 6 | 5422703 | 5425089 |
| VIT_06s0009g00770 | ALDEHYDE OXIDASE 4 | 6 | 12734005 | 12755925 |
| VIT_06s0009g00780 | ABI3-INTERACTING PROTEIN 2 | 6 | 12797100 | 12798180 |
| VIT_06s0009g00790 | SHUGOSHIN 2 | 6 | 12798181 | 12799747 |
| VIT_06s0080g00370 | F-BOX/LRR PLANT PROTEIN | 6 | 21456653 | 21457918 |
| VIT_06s0080g00390 | TRANSMEMBRANE 9 SUPERFAMILY MEMBER<br>5 | 6 | 21484446 | 21490339 |
| VIT_06s0080g00400 | 60S RIBOSOMAL PROTEIN L10A-1-RELATED | 6 | 21491508 | 21499541 |
| VIT_06s0080g00410 | FERREDOXIN-3, CHLOROPLASTIC | 6 | 21518769 | 21522083 |
| VIT_06s0080g00420 | GLYCOLATE OXIDASE 1 | 6 | 21546007 | 21547995 |
| VIT_10s0003g03830 | (S)-2-HYDROXY-ACID OXIDASE GLO1 | 10 | 10085917 | 10090918 |
| VIT_10s0003g03840 | MAJOR FACILITATOR | 10 | 10106763 | 10109792 |
| VIT_10s0003g03850 | PROTEASE DO-LIKE 5, CHLOROPLASTIC | 10 | 10123972 | 10131022 |
| VIT_15s0021g00880 | FERREDOXIN-LIKE PROTEIN | 15 | 10742672 | 10755779 |
| VIT_15s0021g00890 | E3 UBIQUITIN-PROTEIN LIGASE ATL4 | 15 | 10761195 | 10763021 |
| VIT_15s0021g00900 | UNCHARACTERIZED PROTEIN | 15 | 10767831 | 10770627 |
| VIT_15s0021g00910 | DOLICHYL PYROPHOSPHATE<br>GLC1MAN9GLCNAC2 ALPHA-1,3-GLUCOSYLTRANSFERASE-RELATED | 15 | 10773614 | 10778293 |
| VIT_15s0021g00920 | ATP SYNTHASE SUBUNIT DELTA,<br>MITOCHONDRIAL | 15 | 10798637 | 10802728 |
| VIT_15s0021g00930 | TETRATRICOPEPTIDE REPEAT (TPR)-LIKE<br>SUPERFAMILY PROTEIN | 15 | 10803633 | 10807025 |
| VIT_15s0021g01000 | 60S RIBOSOMAL PROTEIN L37A-2 | 15 | 10984273 | 10987176 |
| VIT_15s0021g01010 | HISTONE H3 K4-SPECIFIC<br>METHYLTRANSFERASE SET7/9 FAMILY<br>PROTEIN | 15 | 10988998 | 10993730 |
| VIT_15s0021g01020 | CASEIN KINASE II SUBUNIT BETA-2 | 15 | 11003550 | 11012476 |
| VIT_15s0021g01040 | CYTOCHROME P450 FAMILY PROTEIN | 15 | 11039514 | 11047215 |
| VIT_15s0021g01050 | UNCHARACTERIZED PROTEIN | 15 | 11052160 | 11066622 |
| VIT_15s0046g00990 | HELICASE WITH ZINC FINGER PROTEIN | 15 | 18046425 | 18054747 |
| VIT_15s0046g01000 | TREHALOSE 6-PHOSPHATE PHOSPHATASE | 15 | 18071376 | 18073084 |
| VIT_15s0046g01010 | ATP-DEPENDENT RNA HELICASE MSS116,<br>MITOCHONDRIAL | 15 | 18075965 | 18081383 |
| VIT_15s0046g01020 | NUCLEOTIDYLTRANSFERASE | 15 | 18082315 | 18098641 |
| VIT_17s0000g05750 | ISOCHORISMATE SYNTHASE 2,<br>CHLOROPLASTIC | 17 | 6350085 | 6359530 |
| VIT_17s0000g05760 | YLP MOTIF-CONTAINING PROTEIN 1 | 17 | 6361704 | 6385247 |
| VIT_17s0000g05770 | TRNASE Z TRZ1 | 17 | 6397487 | 6410902 |
| VIT_17s0000g05780 | LANT-SPECIFIC COMPONENT OF THE PRE-RRNA PROCESSING COMPLEX2 | 17 | 6413925 | 6415393 |
| VIT_17s0000g05790 | PLANT CYSTEINE OXIDASE 2 | 17 | 6417479 | 6421337 |
| VIT_17s0000g05800 | DELTA-1-PYRROLINE-5-CARBOXYLATE<br>DEHYDROGENASE 12A1, MITOCHONDRIAL | 17 | 6421832 | 6437350 |
| VIT_17s0000g05810 | WRKY TRANSCRIPTION FACTOR 61-RELATED | 17 | 6447796 | 6451662 |
| VIT_18s0072g01220 | ABC TRANSPORTER G FAMILY MEMBER 25 | 18 | 23880060 | 23887404 |
| VIT_18s0072g01230 | DISEASE RESISTANCE PROTEIN RPP2A | 18 | 23908904 | 23915819 |
| VIT_18s0072g01240 | SULFOTRANSFERASE 12 | 18 | 23934207 | 23938632 |
| VIT_19s0027g00910 | CONTAINING PROTEIN 9, PUTATIVE,<br>EXPRESSED-RELATED | 19 | 20962285 | 20963887 |
| VIT_19s0027g00920 | UNCHARACTERIZED PROTEIN | 19 | 20999186 | 20999990 |

